# The Troyer syndrome protein spartin mediates selective autophagy of lipid droplets

**DOI:** 10.1101/2021.08.18.456894

**Authors:** Jeeyun Chung, Joongkyu Park, Zon Weng Lai, Talley J. Lambert, Ruth C. Richards, Robert V. Farese, Tobias C. Walther

**Affiliations:** Department of Cell Biology, Harvard Medical School, Boston, MA 02115, USA; Department of Molecular Metabolism, Harvard T. H. Chan School of Public Health, Boston, MA, 02115 USA; Harvard T.H. Chan Advanced Multi-omics Platform, Department of Molecular Metabolism, Harvard T. H. Chan School of Public Health, Boston, MA, 02115 USA; Department of Pharmacology, Department of Neurology, Wayne State University School of Medicine, Detroit, MI 48201, USA; Broad Institute of Harvard and MIT, Cambridge, MA, 02124 USA; Howard Hughes Medical Institute, Boston MA 02115 USA

## Abstract

Lipid droplets (LDs) are crucial organelles for energy storage and lipid homeostasis. Autophagy of LDs is an important pathway for their catabolism, but the molecular mechanisms mediating targeting of LDs for degradation by selective autophagy (lipophagy) are unknown. Here we identify spartin as a receptor localizing to LDs and interacting with core autophagy machinery, and we show that spartin is required to deliver LDs to lysosomes for triglyceride (TG) mobilization. Mutations in SPART (encoding spartin) lead to Troyer syndrome, a form of hereditary spastic paraplegia. We find that interfering with spartin function leads to LD and TG accumulation in motor cortex neurons of mice. Our findings thus identify spartin as a lipophagy receptor and suggest that impaired LD turnover may contribute to Troyer syndrome development.

## Main Text

LDs are ubiquitous cytoplasmic organelles that store neutral lipids, such as triglycerides (TGs) or sterol esters, as reservoirs of metabolic fuels and membrane lipid precursors (*1*). As such, they play key roles in metabolism and physiology, and abnormalities in LD biology are increasingly recognized as causes of human disease (*2, 3*).

Key to LD physiology is the ability of cells to mobilize lipids from LDs. This occurs by one of two pathways (*4*). First, lipids can be mobilized from LDs by a series of hydrolytic reactions, known as lipolysis, that is initiated by TG hydrolases such as adipose TG lipase (ATGL), hormone sensitive lipase (HSL), or brain TG lipase (DDHD2) (*5–7*). Alternatively, in a second pathway, selective autophagy delivers LDs to lysosomes, where their lipids are degraded by lysosomal acid lipase, a process also known as lipophagy (*8*). While lipophagy has been recognized for more than a decade and appears to operate in many cell types (*4, 8–11*), the mechanisms and physiological functions of lipophagy remain elusive. One factor limiting our understanding the contributions of lipophagy to lipid homeostasis is that specific factors linking LDs to molecular autophagy machinery are unknown.

Here, we sought to identify the protein machinery that targets LDs for degradation by autophagy (*12*). Selective targeting of LDs to lysosomes predicts the existence of a receptor protein that interacts with both LDs and autophagic machinery, thereby localizing LDs to lysosomal compartments. One candidate for such a receptor is spartin (Spg20). Spartin was previously found to localize to both LDs (*13*) and endosomal compartments (*14, 15*). Spartin contains a ubiquitin-binding region (UBR) (*16, 17*), and ubiquitin binding is a feature of other receptors in selective autophagy, such as p62/SQSTM1 (*18*). We therefore investigated spartin as a potential lipophagy receptor.

To determine whether endogenous spartin localizes to LDs, we genome-engineered human SUM159 cells to express spartin tagged with mScarlet-I from its endogenous genomic locus (Figure S1A). For comparison with a LD marker protein, we also genome-engineered a cell line with endogenously tagged mScarlet-I-PLIN3 and endogenously tagged HaloTag-spartin (Figure S1A). We investigated the localization of spartin by fluorescence microscopy in cells grown in medium containing oleic acid to induce LD accumulation. Spartin localized to LDs stained with BODIPY493/503 or labeled with *LiveDrop,* a fluorescent LD biomarker generated from the hairpin motif of GPAT4 (aa 152-308) (*19*) (Figure 1, A and B and Figure S1, B and C). Spartin localization to LDs was much more apparent 24 hours after oleate treatment than at a 30 min timepoint (Figure 1, A and B). In comparison with PLIN3, which targeted both nascent and mature LDs (Figure S1D), spartin localized primarily to a subset of mature LDs that contained less PLIN3 (Figure 1, C and D). Moreover, the LD population that contained spartin localized preferentially to the cell periphery (Figure 1C and Figure S1B). Collectively, these findings indicate that endogenous spartin localizes to a population of mature LDs, which were often found in the periphery of cells.

**Figure 1.**
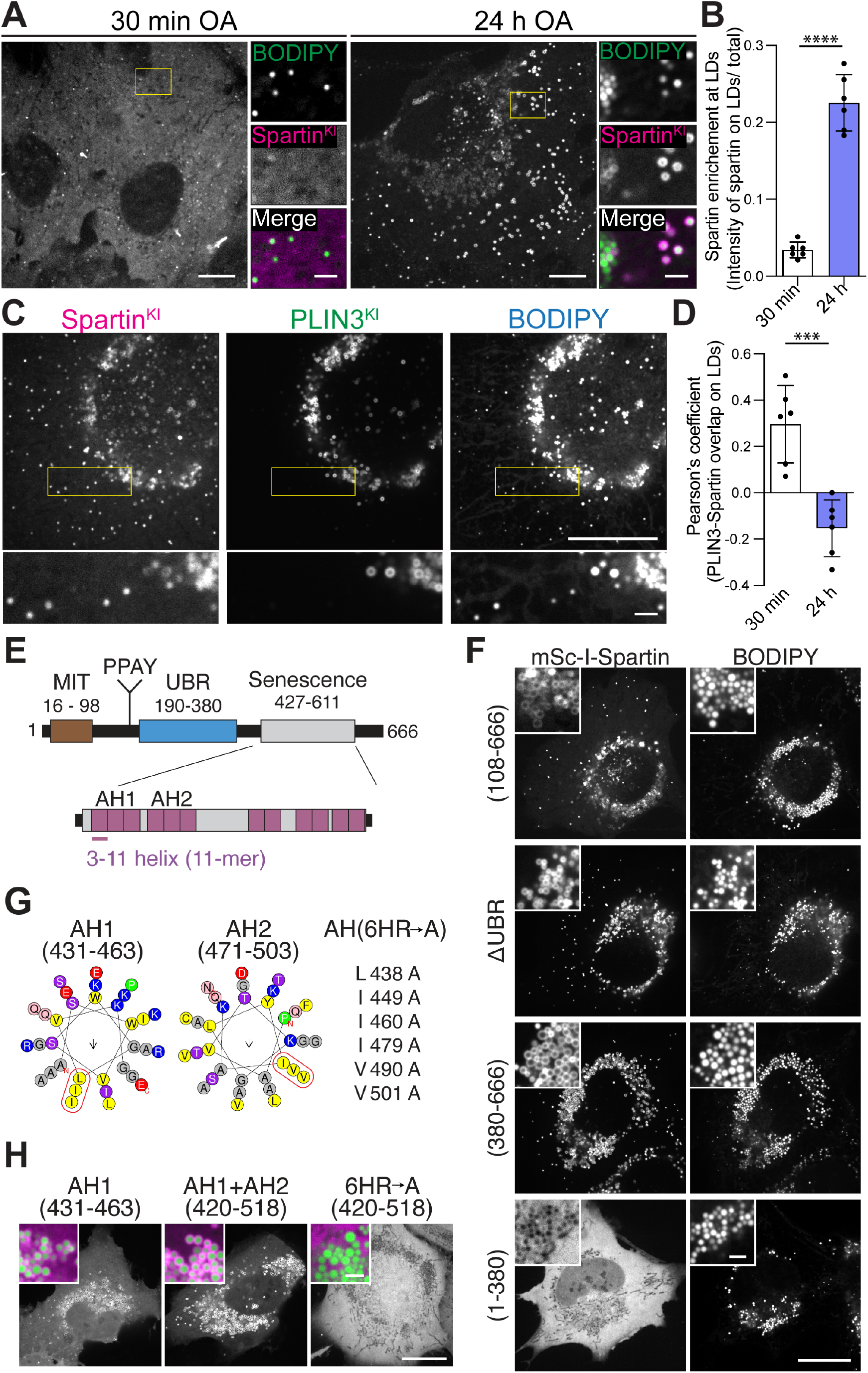
Spartin targets to mature LDs via amphipathic helix repeats in the senescence domain. **(A)** Spartin preferentially targets to mature LDs. Confocal imaging of live SUM159 cells expressing endogenously fluorescent-tagged spartin (with mScarlet-I) at their gene locus. Cells were treated with 0.5 mM oleic acid for 30 mins (left panel) or 24 h (right panel) and stained with BODIPY493/503 prior to image acquisition. Scale bars: full-size, 20 μm; insets, 2 μm. **(B)** Quantification of spartin enrichment at LDs shown in (A). Mean ± SD, n = 6 fields of view, *\*\*\*\*p* < 0.0001, unpaired t-test. **(C)** Spartin and PLIN3 localize to different subpopulations of LDs. Confocal imaging of live SUM159 cells expressing endogenously fluorescent-tagged spartin (with HaloTag) and PLIN3 (with mScarlet-I) at their genomic loci. Cells were treated with 0.5 mM oleic acid for 24 h and stained with BODIPY493/503 prior to image acquisition. HaloTag was pre-labeled with 100 nM JF646. Scale bars: full-size, 20 μm; insets, 2 μm. **(D)** Overlap between PLIN3 and spartin on LDs after 30 min or 24 h 0.5 mM oleic acid treatment was quantified by Pearson’s coefficient analysis. Mean ± SD, n = 6 fields of view, \*\*\**p* < 0.001, unpaired t-test. **(E)** Schematic representation of spartin (top panel) with long AH regions (purple) in the senescence domain (bottom panel). **(F)** The senescence domain of spartin is required for its LD localization. Comparative analysis of localization of spartin truncation mutants (with mScarlet-I tag) when overexpressed in SUM159 cells monitored by confocal microscopy in live cells, treated with 0.5 mM oleic acid for 24 h and stained with BODIPY493/503 prior to image acquisition. Scale bars: full-size, 20 μm; insets, 2 μm. **(G)** Helical wheel plot of 33-mer repeats of AH1 (aa 431-463) and AH2 (aa 471-503) from spartin, plotted as a 3– 11 helix (*36*). The AH(6HR→A) depicting six mutations that were introduced into spartin AH1+AH2 (aa 431-503). **(H)** Spartin AH1 and AH2 are sufficient for LD binding. Confocal imaging of live SUM159 cells expressing mScarlet-I tagged constructs depicted in (G). Cells were treated with 0.5 mM oleic acid for 24 h and stained with BODIPY493/503 prior to image acquisition. Inset showed overlay of spartin (magenta) and LDs (green). Scale bars: full-size, 20 μm; insets, 2 μm.

We next asked how spartin localizes to LDs. Spartin contains three evolutionarily conserved domains—a “microtubule-interacting and trafficking” (MIT) domain (*20*), a “ubiquitin-binding region” (UBR) domain (*16*), and a “plant-related senescence domain (senescence domain)”(*21, 22*) (Figure 1E). Domain truncation analyses of expressed spartin protein revealed that the senescence domain is necessary and sufficient for spartin association with LDs (Figure 1F), in agreement with a recent report (*13, 22*). Analyzing the sequence of the senescence domain for potential regions that may mediate its association with LDs, we detected a series of amphipathic helices (AHs) (Figure 1E, lower panel). AHs are often found in cytosolic proteins that interact with the LD surface, where they integrate into the phospholipid monolayer of LDs (*23–25*). Spartin contains up to 12 sequence stretches with predicted propensity to form 3–11 AHs (11 amino acids per three turns) (Figure 1, E and G), similar to those found in other LD-binding proteins, such as perilipins and ☐-synuclein (*23, 26, 27*). Particularly, the Phyre2 structure prediction revealed that spartin sequences of aa 427–517 shows high sequence and structural similarities to LD targeting AH repeats in α-synuclein (aa 4-93) (*26, 28*). Analyzing these sequences in expression studies, we found that the first extended AH (AH1, containing three 3–11 repeats, aa 431-463) was sufficient to mediate LD binding, and its localization to LDs was enhanced if the second extended AH2 (aa 464-503) was included (Figure 1, G and H). Consistent with the importance of AH1 and AH2 in mediating LD binding, mutating six hydrophobic residues across AH1 and AH2 to alanine abolished association of AH1 and AH2 with LDs (Figure 1, G and H).

For spartin to act as a receptor in lipophagy, it needs to interact not only with LDs but also with the autophagy machinery. To test for this, we co-expressed HA-tagged spartin with 3xFLAG-tagged ATG8-type proteins LC3A, LC3B, LC3C, GABARAP, GABARAPL1 or GABARAPL2 and analyzed potential interactions by assaying for co-immunoprecipitation of the proteins. In these assays, we detected an interaction specifically between spartin and LC3A or LC3C (Figure 2A). Consistent with this finding, spartin-coated LDs co-localized with fluorescently tagged LC3A (Figure 2B).

**Figure 2.**
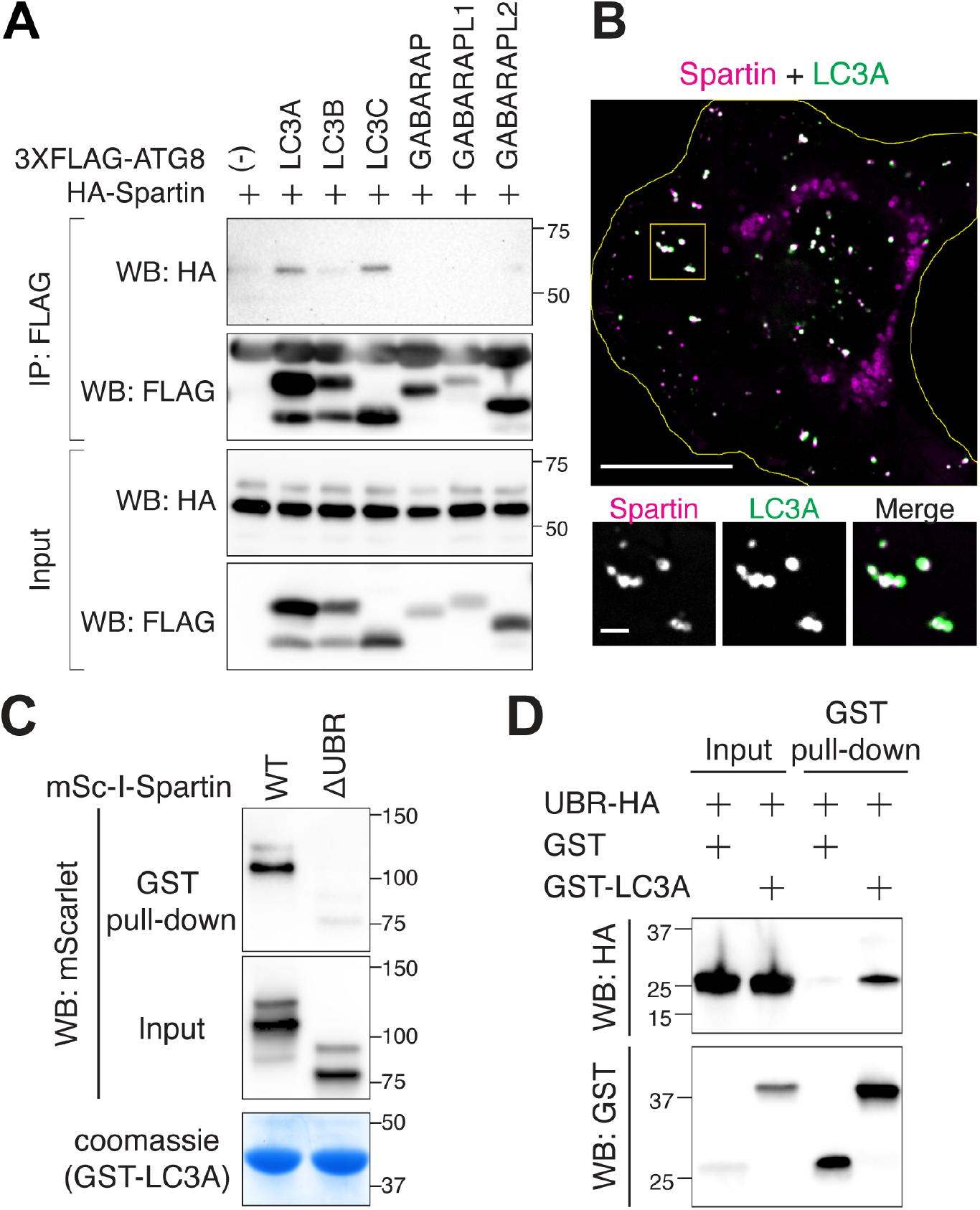
Spartin directly interacts with specific ATG8 family members through its UBR domain. **(A)** Spartin interacts with LC3A and LC3C. HEK293T cells transiently co-expressing HA-spartin together with either 3xFLAG-ATG8 (LC3A, LC3B, LC3C, GABARAP, GABARAPL1, or GABARAPL2) were subjected to immunoprecipitation with anti-FLAG antibody and analyzed by immunoblot with anti-HA and anti-FLAG antibodies. **(B)** Spartin co-localizes with LC3A. Confocal imaging of live SUM159 cells expressing mScarlet-I-spartin full-length and HaloTag-LC3A (pre-labeled with 100 nM JF646). Cells were treated with 0.5 mM oleic acid for 24 h and chased for 3 h after oleic acid withdrawal. Scale bars: full-size, 20 μm; insets, 2 μm. **(C)** The UBR domain is required for the interaction between spartin and LC3A. HEK293T cell lysates transiently expressing mScarlet-I-spartin WT or ΔUBR were incubated with recombinant GST-LC3A, followed by GST pull-down and detected with anti-mScarlet/mCherry antibody. Cells were treated will 0.5 mM OA for 24 h prior to lysate preparation. **(D)** Recombinant UBR domain directly interacts with recombinant LC3A. Recombinant spartin-UBR-HA was incubated with recombinant GST or GST-LC3A, followed by GST pull-down and detected with anti-GST and anti-HA antibodies.

To map the domain of spartin that is required for the interaction with LC3A, we performed GST-pull-down analyses of recombinant LC3A with HEK293T cell lysates expressing full-length spartin or spartin lacking the UBR domain. We found that the UBR domain is required for spartin’s association with recombinant LC3A (Figure 2C and Figure S2, A to C). To corroborate these results, we purified recombinantly produced LC3A and the UBR domain of spartin (Figure S2D) to assay for direct interactions between the proteins. By performing a pull-down assay with LC3A, we found it directly interacted with the UBR domain (Figure 2D). Interactions between autophagy receptors and LC3-type proteins are often, but not always, mediated by a short LC3-interacting region (LIR) motif (*29, 30*). The spartin UBR contains at least two amino acid stretches with similarity to the LIR consensus motif (aa 194-197 and aa 276-279). However, neither of these sequences was required for spartin’s interaction with LC3A (not shown), indicating that binding between spartin and LC3A likely occurs in a different, possibly non-canonical manner.

If spartin participates in the lysosomal degradation of LDs, then spartin-marked LDs should co-localize with lysosomes. Fluorescence microscopy revealed that a subset of *LiveDrop*-labeled LDs co-localized with lysosomes, marked by expression of fluorescently tagged LAMP1 (Figure 3A). This colocalization between the two organelles required spartin, inasmuch as it was largely reduced in spartin knockout (KO) cells (Figure 3, A and B and Figure S3, A and B). Additionally, we found that co-localization of spartin-marked LDs with LAMP1 was dependent on spartin’s UBR domain (Figure 3C). To test whether these co-localized organelles are dependent on autophagy machinery, we assessed for spartin and LAMP1 colocalization in the presence of knockouts of either ATG5, ATG7, or FIP200 (Figure S3C). We found that each of these proteins was required for the association of LDs and lysosomes (Figure 3, D and E). Additionally, we found no differences between wild-type and spartin KO cells in an assay monitoring LC3 lipidation (conversion of LC3-I to LC3-II) during nutrient starvation (Figure S3D), indicating spartin is not required for autophagy in general. Collectively, these results suggest that spartin interacts through the senescence domain with LDs to recruit core autophagy machinery via its UBR, and thus delivers the organelle to lysosomes.

**Figure 3.**
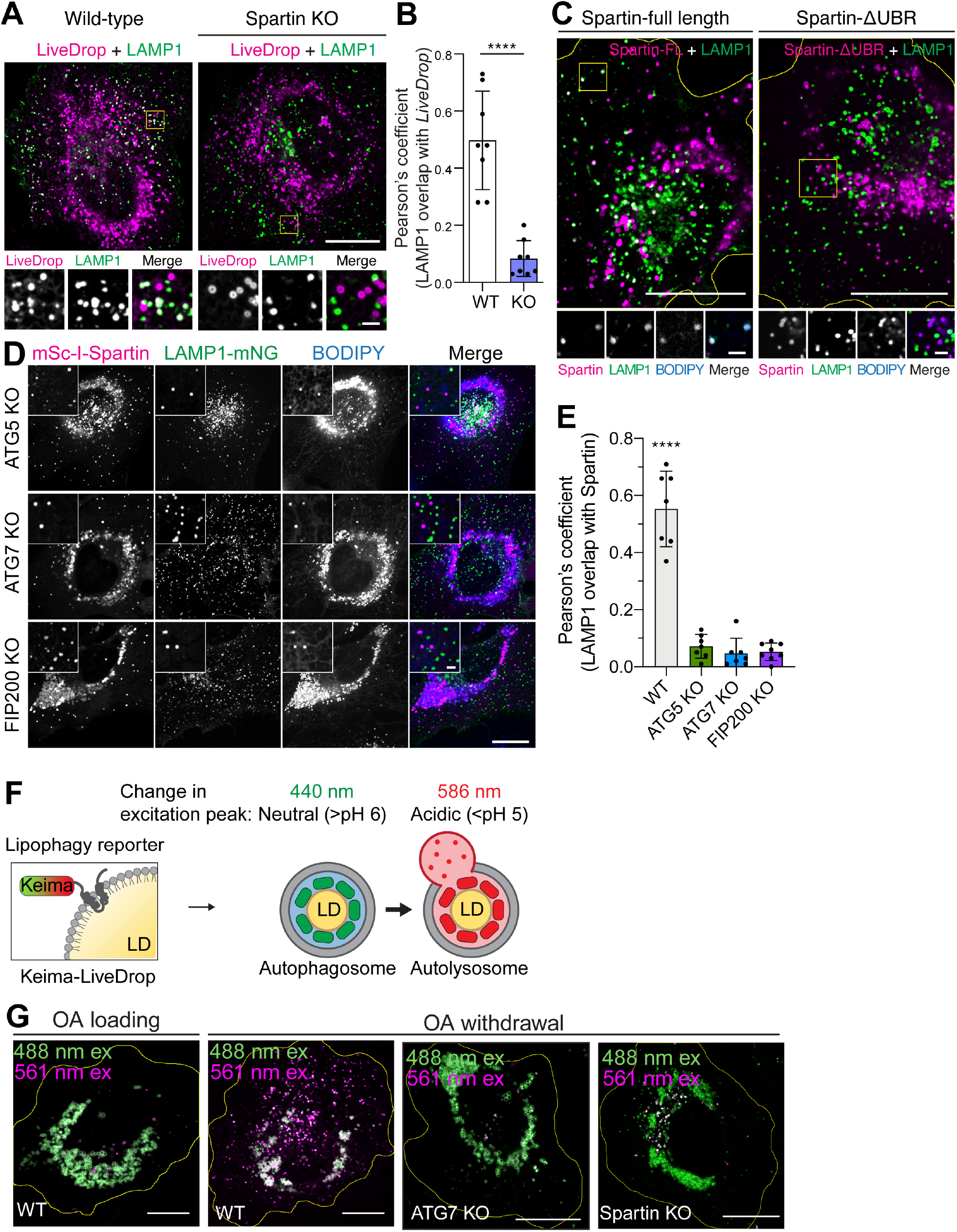
Spartin mediates autophagy-dependent LD delivery to lysosomes. (**A**) Spartin deficiency impairs LD targeting to lysosomes. Confocal imaging of live SUM159 wild-type and spartin KO cells transiently co-expressing HaloTag-*LiveDrop* (pre-labeled with 100 nM JF549) and LAMP1-mNG. Cells were treated with 0.5 mM oleic acid for 24 h and chased for 3 h after oleic acid withdrawal. Scale bars: full-size, 20 μm; insets, 2 μm. **(B)** Overlap between *LiveDrop* and LAMP1 in cell periphery shown in (A) was quantified by Pearson’s coefficient analysis. Mean ± SD, n = 8 cell periphery region of interest from 4 cells, \*\*\*\**p* < 0.001, unpaired t-test. **(C)** UBR domain mediates associations between spartin-coated LDs and lysosomes. Confocal imaging of live SUM159 spartin KO cells transiently co-expressing mScarlet-I-spartin full length or ΔUBR together with LAMP1-mNG. Cells were treated with 0.5 mM oleic acid for 24 h and chased for 3 h after oleic acid withdrawal. Scale bars: full-size, 20 μm; insets, 2 μm. **(D)** Delivery of spartin-coated LDs to lysosomes is autophagy-dependent. Co-localization analyses between transiently overexpressed mScarlet-I-spartin and LAMP1-mNG in SUM159 cells lacking ATG5, ATG7, or FIP200. Scale bars: full-size, 20 μm; insets, 2 μm. **(E)** Overlaps between spartin and LAMP1 in cell periphery shown in (D) were quantified by Pearson’s coefficient analysis. Mean ± SD, n = 7 cell periphery crops from 4 cells, *p* < 0.0001, one-way ANOVA followed by post hoc Dunnett’s multiple comparisons. **(F)** Schematic illustration showing how Keima-*LiveDrop* works. **(G)** Excitation spectrum converstion of Keima in varius conditions. Overlay live-cell confocal images of Keima-*LiveDrop* acquired by 488 nm (green) and 561 nm (magenta) excitations expressed in SUM159 wild-type (first two panels), ATG7 KO (third panel from left), or spartin KO (far right panel). Scale bars: 10 μm.

Our results prompted us to test directly whether spartin was required for LD degradation by lipophagy in the lysosome. To test this, we developed a lipophagy reporter system based on the Keima-fluorophore. This fluorescent protein is sensitive to pH and shifts its excitation spectrum upon trafficking to the lysosome (pH of ∼4.5)(*31*). We fused Keima to the N-terminus of *LiveDrop* to generate a fluorescence reporter that selectively shifts fluorescence depending on whether LDs are located in the cytoplasm versus the lysosome (Figure 3F). Indeed, withdrawal of oleic acid from the cell culture media led a shift in Keima fluorescence consistent with LDs being engulfed by lysosomes (Figure 3G, first left two panels). The transition of *LiveDrop* Keima-fluorescence due to lysosomal localization was dependent on an intact autophagy pathway as it was abolished in ATG7 KO cells (Figure 3G, third left panel). Similarly, deletion of spartin also prevented this change in Keima emission spectrum, indicating spartin is required for lipophagy of this reporter (Figure 3G, the most right panel).

Spartin mediating degradation of LDs by selective autophagy predicts an increase in stored lipids in cells lacking spartin. To test this, we measured TG synthesis, accumulation, and turnover in wild-type and spartin KO cells. After 24 h of oleate addition, spartin KO cells accumulated more TG than wild-type cells (Figure 4A). We found no differences between wild-type and spartin KO cells in cellular TG synthesis assays (Figure 4B) or in short-term (30 min) LD formation assays (Figure S4), indicating that TG accumulation in spartin KO cells was not likely due to altered rates of LD biogenesis. To assess the possibility that spartin affected catabolism of LDs, we utilized an assay in which we first incubated SUM159 cells in medium containing fatty acids, resulting in the formation of abundant LDs, and then withdrew lipids to stimulate LD turnover. In this assay, visualizing LDs by fluorescence microscopy revealed that spartin KO cells had impaired LD degradation, with many more and larger LDs remaining after 6 h of fatty acid removal from the cell culture medium (Figure 4C). Additionally, pulse-chase assays using radio-labeled fatty acids to trace degradation of TG showed that spartin KO cells had ∼48% lower rates of TG degradation during three hours after fatty acid withdrawal (Figure 4D). A similar impairment of TG degradation was found in cells lacking a core component of the autophagy machinery (ATG7) (Figure 4D). The degree of impairment in TG degradation for spartin KO cells was comparable (∼35–40% reduction) in wild-type cells and ATGL knockdown cells (Figure 4E and Figure S5), indicating that the TG degradation defect was independent of the major TG lipase. Taken together, these data are consistent with a model in which, under conditions of lipid withdrawal, spartin deficiency impairs LD turnover by lipophagy.

**Figure 4.**
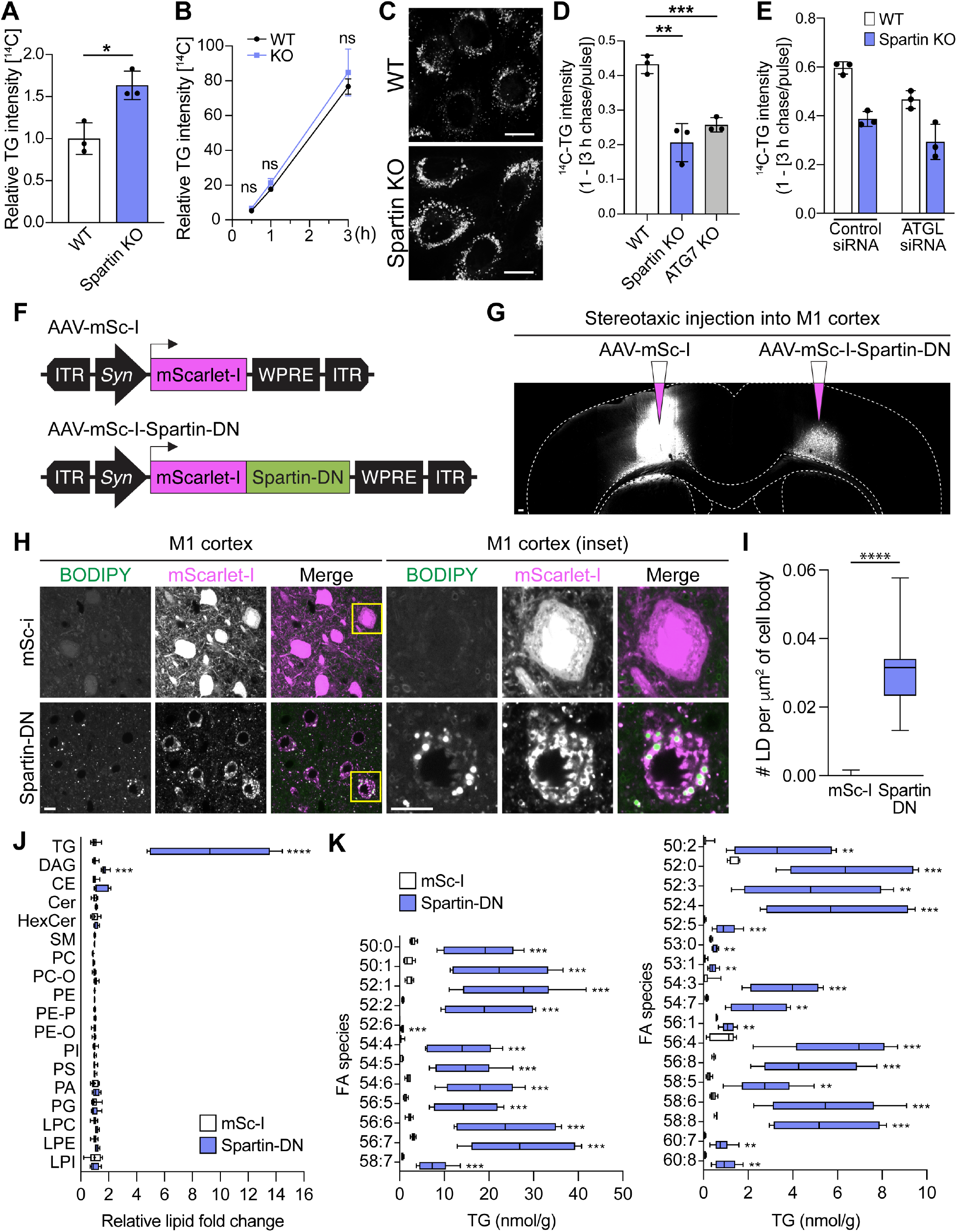
Interfering with spartin function impairs TG turnover in cells and leads to TG and LD accumulation in neurons of the mouse motor cortex. **(A and B)** Spartin KO leads to impairment in TG turnover. WT and spartin KO SUM159 cells were pulse-labeled with [^14^C]-OA, and incorporation into TG was measured after 0.5 mM oleic acid treatment for 24 h (A), 0.5–3 h (B). Values were calculated relative to wild-type (A) and spartin KO cells’ highest value at 3 h. Mean ± SD, n = 3 independent experiments, \**p* < 0.05, n.s. not significant, unpaired t-test (A) and two-way ANOVA with repeated measurements (B). **(C)** Spartin KO results in deficient LD turnover. Confocal live-cell imaging showing LDs (BODIPY493/503 stained) in WT or spartin KO SUM159 cells. Cells were treated with 0.5 mM oleic acid for 24 h and chased for 6 h after oleic acid withdrawal. Scale bar: 20 μm. **(D** and **E)** ATGL-independent reduction of TG degradation in spartin KO. WT, spartin KO, or ATG7 KO SUM159 cells were pulse-labeled with [^14^C]-OA, and incorporation into TG was measured after 0.5 mM oleic acid treatment for 24 h and subsequent 3 h oleic acid withdrawal. ATGL was knocked down in WT and spartin KO cells for 48 h prior to [^14^C]-OA labeling (E). Mean ± SD, n = 3 independent experiments, \*\**p* < 0.01 and \*\*\**p* < 0.001, unpaired t-tests. **(F)** Schematic diagram illustrating AAVs expressing either mScarlet-I as a control (mSc-I) or a dominant-negative form of spartin (spartin-DN) under a pan-neuronal *synapsin* promoter (*Syn*). **(G)** Expression of mScarlet-I and mScarlet-I-fused spartin-DN in the mouse motor cortex. AAV-mSc-I or AAV-mSc-I-spartin-DN was stereotaxically injected into each hemisphere of the M1 motor cortex in 7-8-week-old wild-type mice. Scale bar: 100 μm. **(H** and **I)** Robust accumulation of BODIPY493/503 in spartin-DN-expressing neurons of the mouse M1 motor cortex. Representative images of the AAV-infected M1 cortex slices stained with BODIPY493/503 10-11 days after AAV injection (H). Scale bar: 10 μm. Quantification of LD numbers per μm^2^ of the cell body of the M1 cortex slices (I). Mean ± SD, n = 9 fields of view from n = 3 mice, *\*\*\*\*p* < 0.0001, unpaired t-test. **(J** and **K)** Significant elevation of TGs and DAG in neurons where spartin function was disrupted. Lipidomic profiles of the AAV-infected M1 cortices. Lipids were extracted from tissues using an MTBE method and analyzed by LC–MS in both positive and negative ion mode as described in Materials and Methods. Relative fold changes of different lipid classes (J). LC–MS analysis verified widespread elevations in TGs from spartin-DN-expressing neurons of the M1 cortex (Spartin-DN) compared to a control (mSc-I) (K). Mean ± SD, n = 6 mice, *\*\*p* < 0.01, \*\*\**p* < 0.001, *\*\*\*\*p* < 0.0001, unpaired t-tests.

In humans, loss of spartin function due to *SPART* mutations leads to Troyer syndrome, a complex hereditary spastic paraplegia with degeneration of neurons (*32, 33*). Our findings led us to consider whether impaired spartin function in neurons compromises lipophagy in this cell type. Neurons normally do not contain large amounts of TGs or LDs, but deficiency of the neuronal TG lipase DDHD2/Spg54, which is also associated with development of hereditary spastic paraplegia, causes TG accumulation in neurons (*7, 34*), providing evidence that these cells can synthesize and degrade TGs.

Immunohistochemistry staining of endogenous spartin revealed that spartin is ubiquitously expressed in murine brain and widely detectable in neurons (Figure S6). To enable testing how impairing spartin function affects LDs in neurons, we identified a dominant-negative fragment of spartin (aa 380-666, spartin-DN) that binds to LDs but does not engage the autophagic machinery; expression of spartin-DN in the culture cells triggered accumulation of LDs (Figure S7). We injected into one hemisphere of the mouse M1 motor cortex an adeno-associated virus (AAV) expressing mScarlet-I-spartin-DN under the *synapsin* promoter, leading to unilateral interference with spartin function (Figure 4, F and G). As a control, we injected an AAV expressing mScarlet-I into the contralateral hemisphere (Figure 4, F and G). Fluorescence microscopy showed that this resulted in robust expression of mScarlet-I or mScarlet-I-spartin-DN in neurons (Figure 4G). Co-staining of these neurons with the LD probe BODIPY493/503 showed a marked increase of LDs specifically in cells where spartin function was inhibited (Figure 4, H and I). Consistent with an increase in LDs, spartin-DN-expressing cortical neurons showed ∼10-fold increased levels of total TGs, with a marked increase across the detected TG species, compared with mScarlet-I controls (Figure 4, J and K and Figure S8). DAG levels were also increased, but sphingolipid and phospholipid levels were similar in the two regions (Figure 4J and Figure S8). These data indicate that spartin has an important function in mediating TG turnover in neurons of the mouse motor cortex.

Taken together, our data indicate that spartin links LDs to the core autophagy machinery as a lipophagy receptor. This model is supported by spartin’s ability to bind LDs as well as LC3A/C, core components of the autophagy machinery and is consistent with the effect of spartin deficiency in cells, where it leads to accumulation of LDs and TGs. Whether additional lipophagy receptors exist for different cell types or different metabolic conditions is unknown, but remains an area of active investigation (*35*). The current findings show that the spartin-mediated lipophagy pathway may be important in neurons where its impairment leads to accumulation of LDs. Inasmuch as deficiency of either spartin or DDHD2 result in neuronal TG accumulation by different mechanisms, yet both lead to hereditary spastic paraplegia, the available data suggest that impaired TG turnover in neurons may be directly linked to neurodegeneration in humans.

## Acknowledgments

We thank members of Farese & Walther laboratory and S. Kim for helpful discussions, W. Harper for helpful discussion and for sharing of reagents, I. Dikic and T. Kirchhausen for sharing reagents, X. Wu for advice on *in vitro* GST-pull down analysis and for kindly sharing p12xHis-SUMO-plasmid. Other gifts of reagents are acknowledged in the Materials and Methods section. This work was supported by NIH R01GM124348 (to R.V.F.), NIH R01GM097194 (to T.C.W), and NIH R21AG068423 (to J.P.). J.C is a fellow of the Damon Runyon Cancer Research Foundation. T.C.W. is a Howard Hughes Medical Institute Investigator.

## Author contributions

J.C., R.V.F., and T.C.W. conceived the project; J.C., J.P., T.C.W., and R.V.F. designed the experiments; and J.C. performed and analyzed most of the experiments. J.P. performed virus production, injections, and mouse brain histology work. Z.W.L. performed lipidomics analyses. T.J.L. wrote scripts for imaging analyses, and T.J.L. and J.C. analyzed imaging data together. R.C.R helped J.C. for molecular cloning experiments. J.C., T.C.W., and R.V.F. wrote the manuscript. All authors discussed the results and contributed to the manuscript.

## Data and materials availability

All data are available in the main text or the supplementary materials. Further information and requests for resources and reagents should be directed to and will be fulfilled by the T.C.W. or R.V.F. Requests will be handled according to the Harvard T. H. Chan School of Public Health policies regarding MTA and related matters.

---

**Figure S1.**
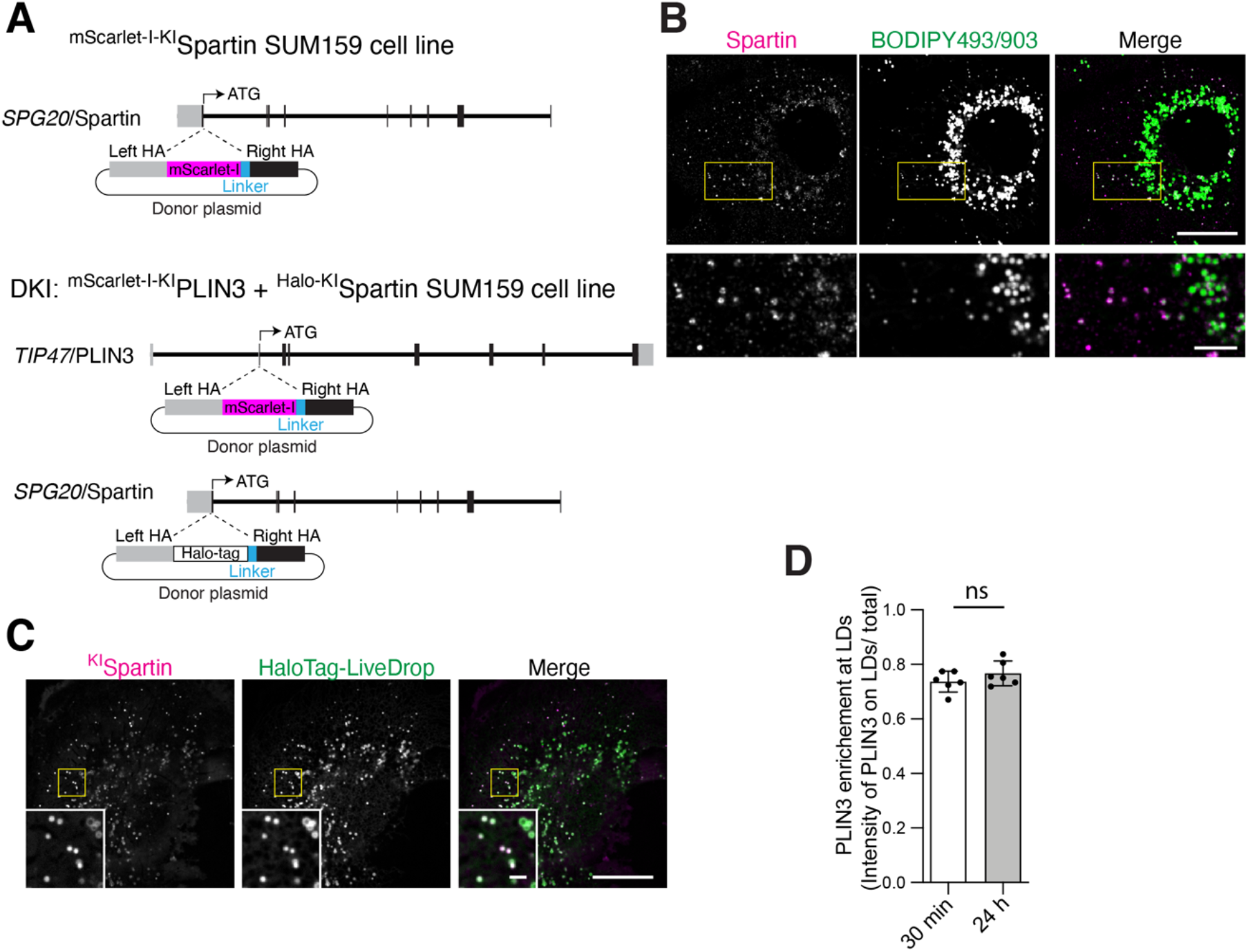
Endogenous spartin targets to LDs at cell periphery. **(A)** Strategy for the endogenous tagging of *SPG20/spartin* and *TIP47/PLIN3*. Individual rectangular bar represents exon (gray color, untranslated region; black color, translated region). A donor plasmid was co-transfected with appropriate gRNA/Cas9 plasmid to induce homologous recombination of mScarlet-I or HaloTag into upstream of *SPG20/spartin* or *TIP47*/PLIN3. **(B)** Confocal imaging of fixed SUM159 cells stained with endogenous spartin antibody and BODIPY493/503. Cells were treated with 0.5 mM oleic acid for 24 h. Scale bars: full-size, 20 μm; insets, 2 μm. **(C)** Confocal imaging of live SUM159 cells expressing endogenously fluorescent-tagged spartin (with mScarlet-I) at their gene locus. HaloTag-*LiveDrop* was transiently expressed and stained with 100 nM JF646. Cells were treated with 0.5 mM oleic acid for 24 h. Scale bars: full-size, 20 μm; insets, 2 μm. **(D)** Quantification of PLIN3 enrichment at LDs in SUM159 cells expressing endogenously fluorescent-tagged spartin (with HaloTag) and PLIN3 (mScarlet-I). Cells were pre-incubated with 0.5 mM oleic acid for 30 min or 24 h prior to confocal live-cell microscopy. Mean ± SD, n = 6 fields of view, n.s. not significant, unpaired t-test.

**Figure S2.**
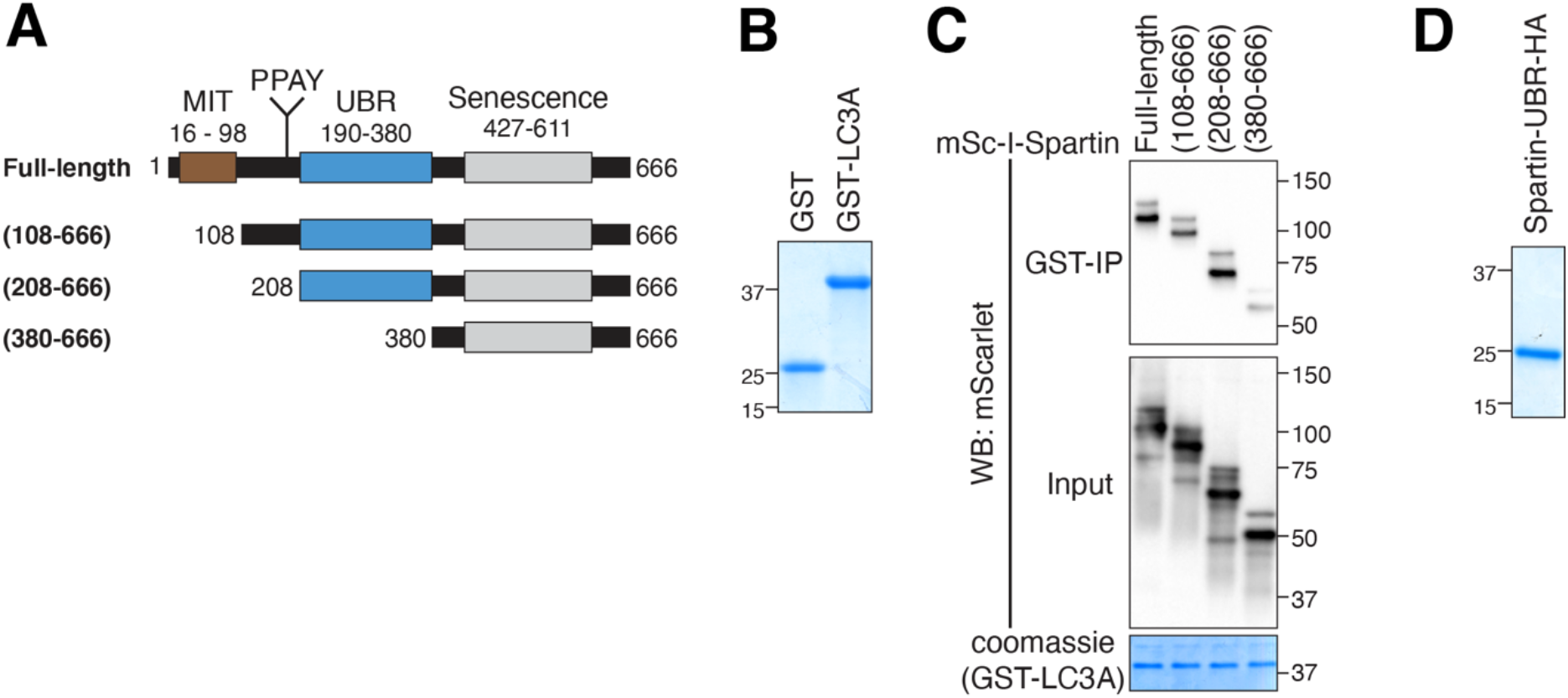
Domain mapping of spartin for LC3A interaction. **(A)** Schematic representation of spartin truncation mutants used in GST pull-down analysis. **(B)** SDS-PAGE analysis of recombinant GST and GST-LC3A purified from bacteria (after size-exclusion column). Proteins were visualized by Coomassie blue staining. **(C)** HEK293T cell lysates transiently expressing mScarlet-I-tagged spartin domain truncation mutants depicted in (A) were incubated with recombinant GST-LC3A, followed by GST pull-down and detected with anti-mScarlet/mCherry antibody. Cells were treated will 0.5 mM OA for 24 h prior to lysate preparation. **(D)** SDS-PAGE analysis of recombinant spartin-UBR-HA purified from bacteria. 12XHis-SUMO tag was removed by SUMO protease and the protein is further purified by size-exclusion column. Protein was visualized by Coomassie blue staining.

**Figure S3.**
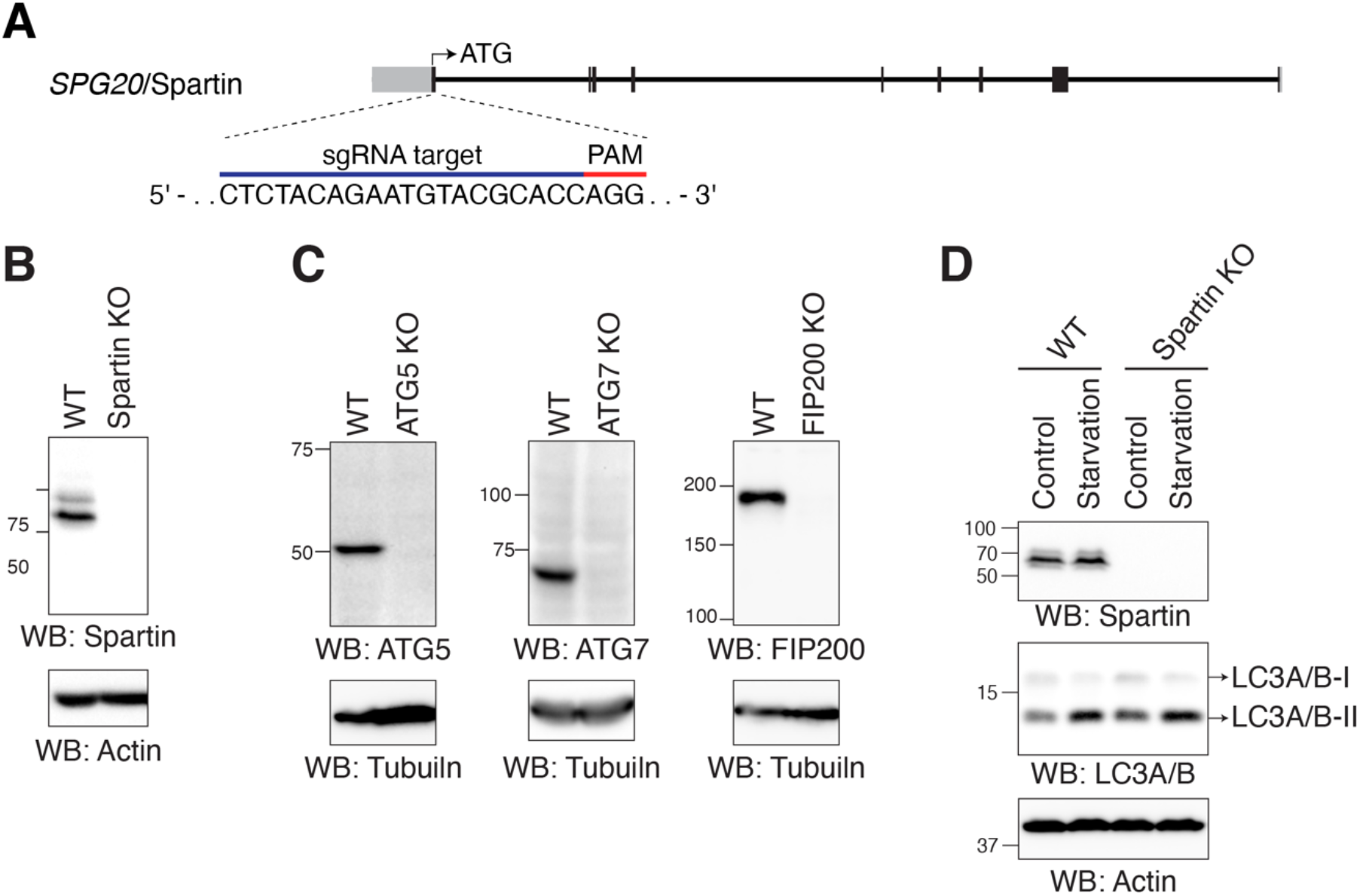
Establishment of cell lines lacking spartin or autophagy machineries. **(A)** Strategy for the knockout of *SPG20/spartin*. Individual rectangular bar represents exon (gray color, untranslated region; black color, translated region). A sgRNA target sequence is shown with PAM sequence. **(B** and **C)** Immunoblot analyses of SUM159 cells wild-type and cells lacking spartin (B) or ATG5, ATG7, or FIP200 (C). **(D)** Immunoblot analyses of SUM159 wild-type and spartin KO cells. Starvation samples were incubated in serum free and low glucose media (1 g/L) for 3 h prior to lysate preparation.

**Figure S4.**
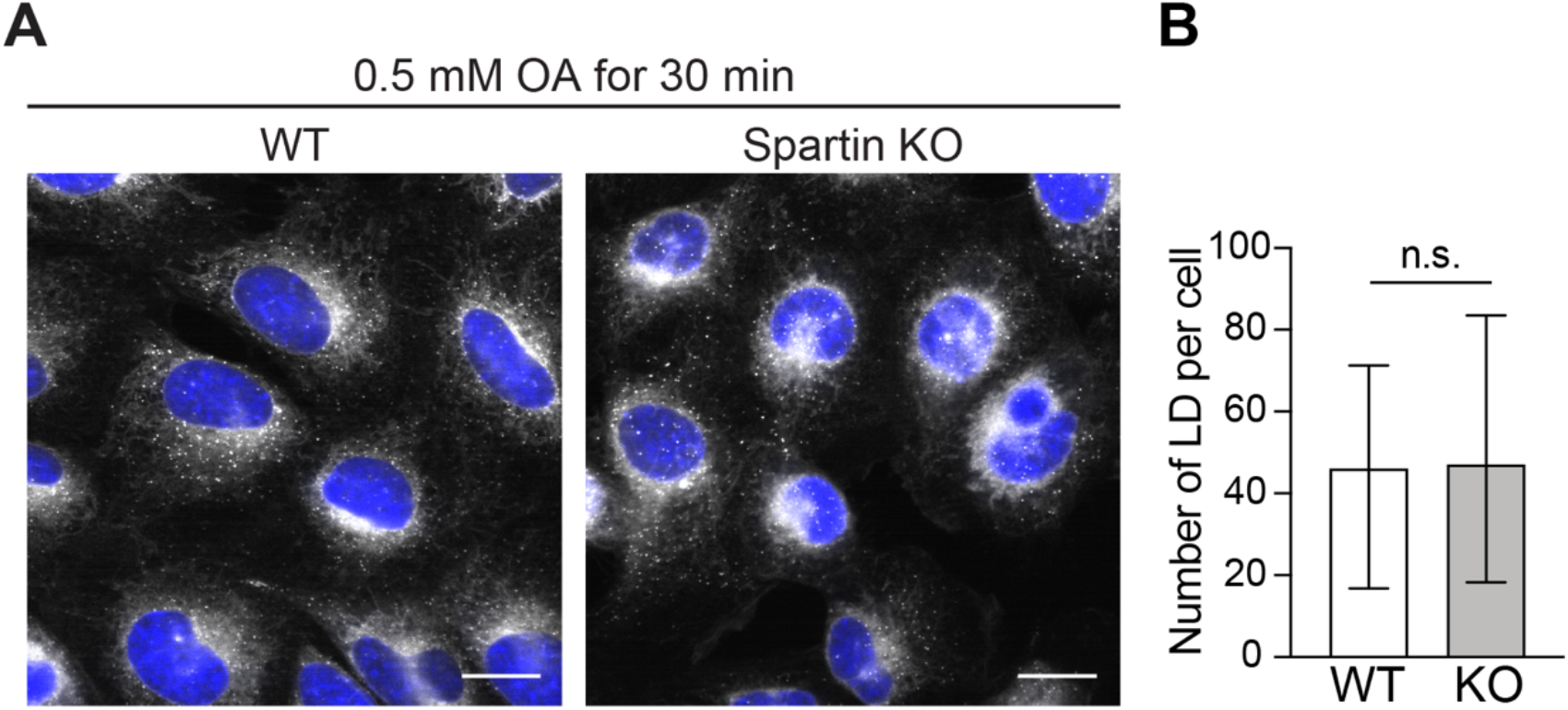
LD biogenesis is not impaired in spartin deficiency. **(A)** Confocal imaging of fixed SUM159 wild-type and spartin KO cells stained with BODIPY493/503 and Hoechst 33342. Cells were treated with 0.5 mM OA for 30 mins prior to fixation. Scale bar: 20 μm. **(B)** Quantification of number of LDs per cell in wild-type and spartin KO SUM159 cells shown in (A) The images were taken on high-throughput confocal microscope. n > 195 cells, median with interquartile range, n.s. not significant, unpaired Welch’s t-test.

**Figure S5.**
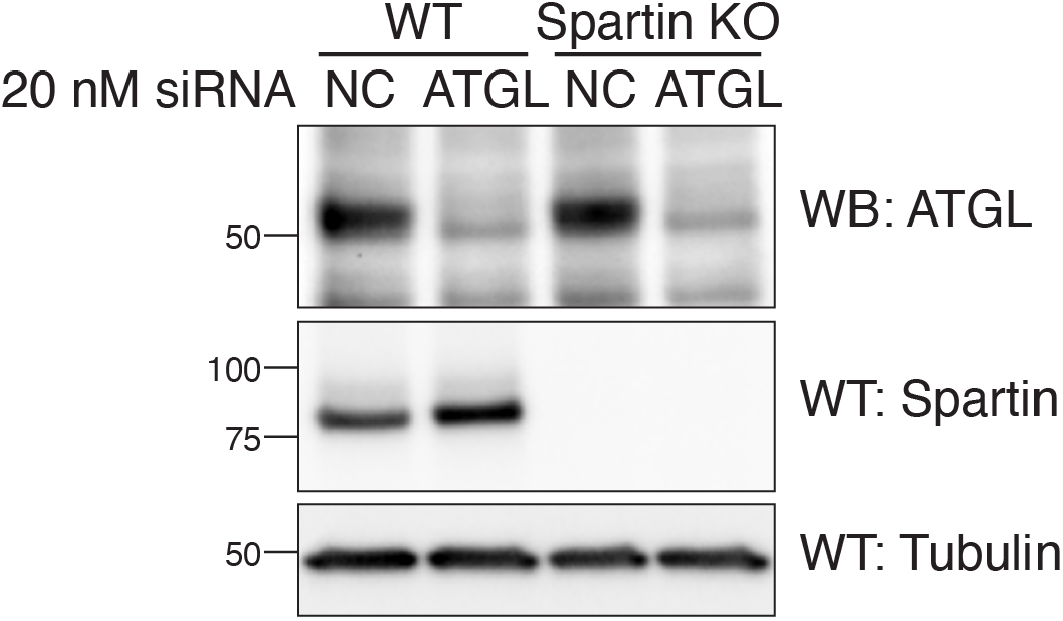
**Validation of ATGL knock-down efficiency** Immunoblot analyses of SUM159 wild-type and spartin KO cells. Cells were treated with 20 nM negative control (NC) siRNA or ATGL siRNA for 72 h prior to lysate preparation.

**Figure S6.**
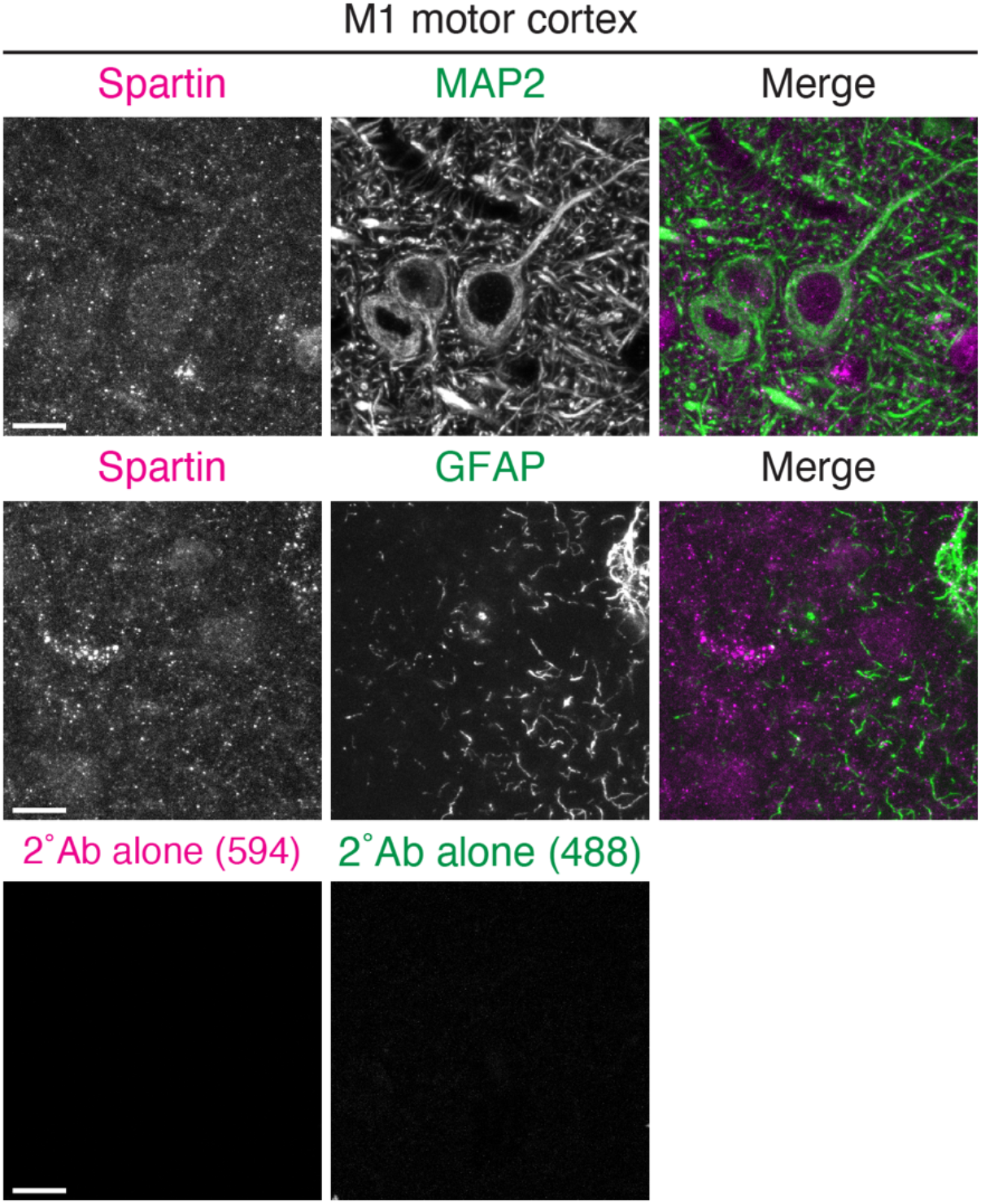
**Spartin endogenously expresses in neurons of the mouse motor cortex.** Confocal micrographs of the M1 motor cortex slices from 9-week-old wild-type mice. PFA-fixed sagittal sections were co-stained with anti-Spartin (magenta) and anti-MAP2 (a neuronal marker; green in the top panels) or anti-GFAP (an astrocyte marker; green in the second panels). Indicate that Spartin signals locate on neuronal cell bodies and projections (top panels). Slices stained with secondary antibodies alone (Alexa Fluor 488 and 594; the bottom panels) were used as background signal controls. Scale bar: 10 μm.

**Figure S7.**
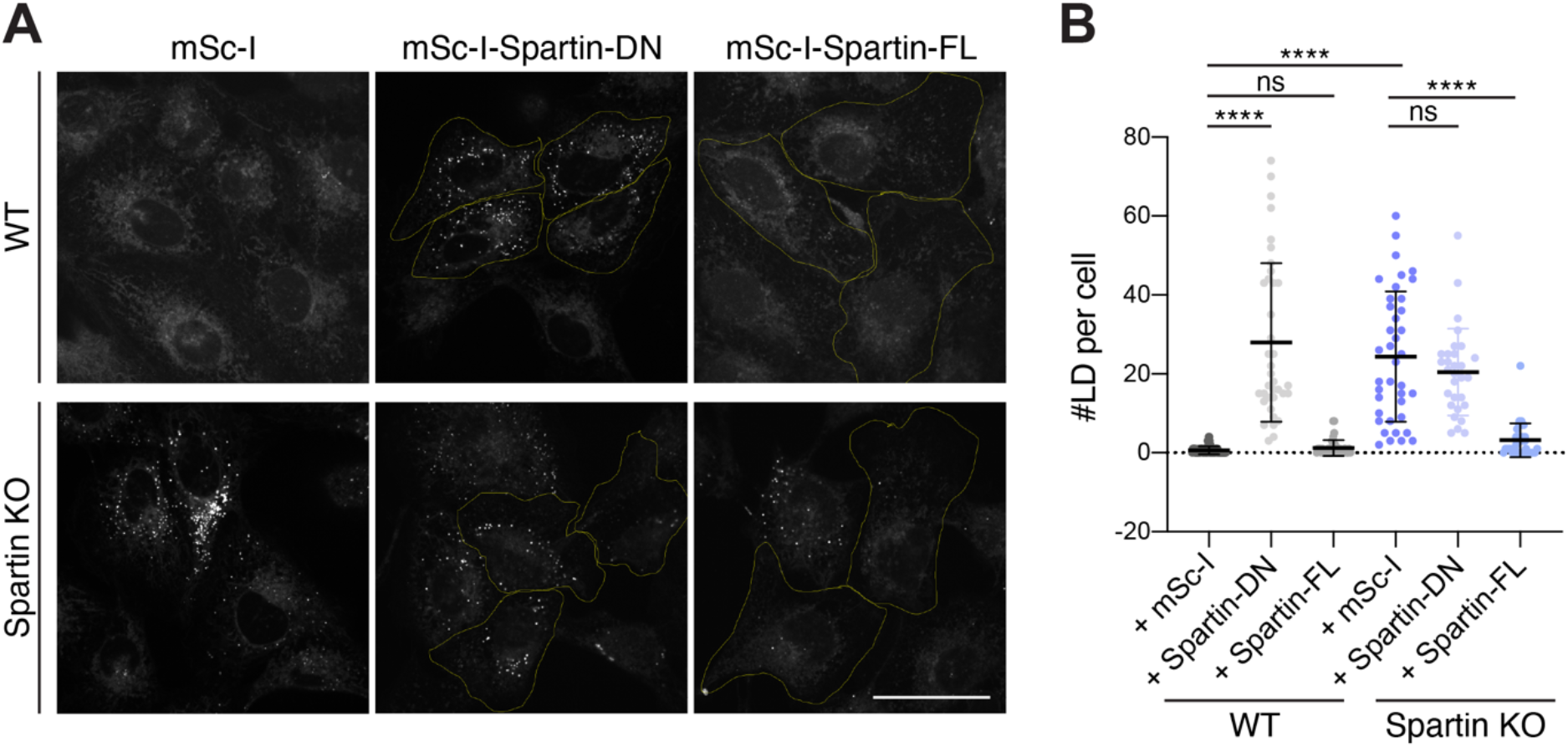
Spartin-DN impairs LD turnover in SUM159 cells. **(A)** Confocal imaging of live SUM159 wild-type (top panel) and spartin KO (bottom panel) cells transiently expressing mScarlet-I-tagged spartin-DN or spartin full-length (spartin-FL) proteins. LDs were stained with BODIPY493/503. Transfected cells were marked with yellow outline. Scale bars: 50 μm. **(B)** Quantification of LD numbers per cell shown in (A). Mean ± SD, n > 30 cells, n.s., not significant, \*\*\*\**p* < 0.0001, unpaired t-tests.

**Figure S8.**
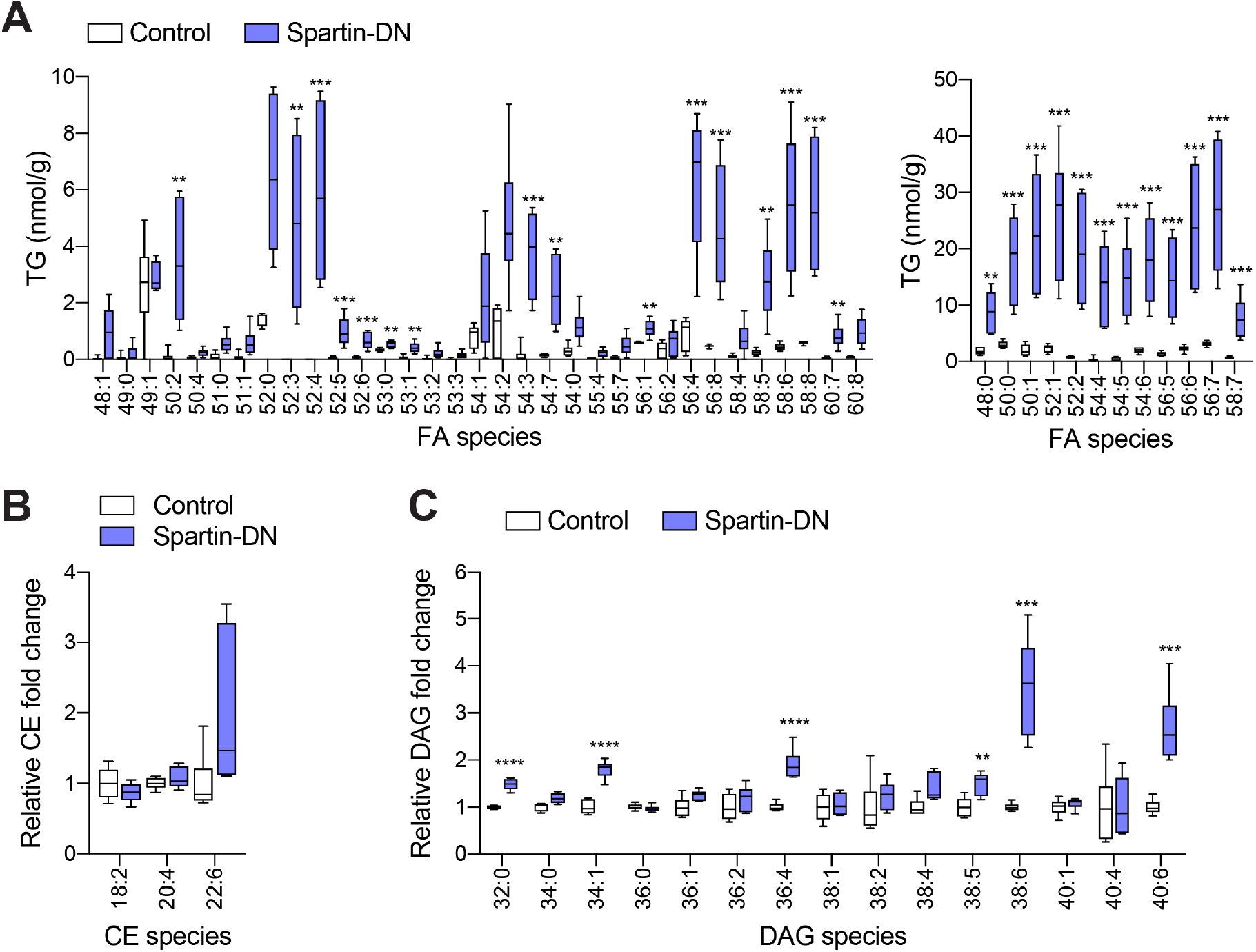
Major lipid class changes in the spartin-DN-infected M1 cortices. **(A to C)** Lipidomic profiles of the AAV-infected M1 cortices. Lipids were extracted from tissues using an MTBE method and analyzed by LC–MSMS positive ion mode. Absolute amounts of detected TG species, normalized by tissue weight (A). Relative fold changes of CE (B) and DAG (C) lipid classes. Mean ± SD, n = 6 mice, \*\**p* < 0.01, \*\*\**p* < 0.001, \*\*\*\**p* < 0.0001, unpaired t-tests.

## Materials and Methods

### Cell culture

SUM159 breast cancer cells were obtained from the laboratory of Dr. Tomas Kirchhausen (Harvard Medical School) and were maintained in DMEM/F-12 GlutaMAX (Life Technologies) supplemented with 5 μg/ml insulin (Cell Applications), 1 μg/ml hydrocortisone (Sigma), 5% FBS (Life Technologies 10082147; Thermo Fisher), 50 μg/ml streptomycin, and 50 U/ml penicillin. Where noted, cells were incubated with media containing 0.5 mM oleic acid complexed with essentially fatty acid–free BSA. For immunoprecipitation analyses, HEK293T cells (ATCC) were used. HEK293T cells were cultured in DMEM (Gibco) supplemented with 10% FBS (Life Technologies), 50 μg/ml streptomycin, and 50 U/ml penicillin.

### Special reagents and antibodies

Janelia Fluor^®^ dyes with HaloTag^®^ (JF549 and JF646) (*37*) were kind gifts of Dr. Luke Lavis (Janelia Research Campus). BODIPY493/503, HCS Lipid TOX Deep Red Neutral Lipid Stain, and puromycin were purchased from Thermo Fisher Scientific. Oleic acid was purchased from Millipore-Sigma. Oleic acid [1-^14^C] was purchased from American Radiolabeled Chemicals.

Primary antibodies used in this study were: rabbit polyclonal anti-SPG20/spartin (Proteintech), rabbit polyclonal anti-FIP200/RB1CC1 (Proteintech), mouse monoclonal anti-FLAG (Millipore-Sigma), anti-FLAG® M2 affinity gel (Millipore-Sigma), rat monoclonal anti-HA (Millipore-Sigma), mouse monoclonal anti-α-tubulin (Millipore-Sigma), rabbit polyclonal anti-mCherry (for detection of mScarlet-I, Abcam), rabbit polyclonal anti-LC3A/B (Cell Signaling Technology), mouse monoclonal anti-actin (Cell Signaling Technology), rabbit polyclonal anti-ATGL (Cell Signaling Technology), rabbit monoclonal anti-ATG5 (Cell Signaling Technology), rabbit monoclonal anti-ATG7 (Cell Signaling Technology), chicken polyclonal anti-MAP2 (Synaptic Systems), rat polyclonal anti-GFAP (Thermo Fisher Scientific), rabbit polyclonal anti-GST (Thermo Fisher Scientific). HRP-conjugated secondary antibodies against mouse and rabbit were from Santa Cruz Biotechnology. Fluorescent secondary antibodies were used in this study were purchased from Thermo Fisher Scientific including DyLight 488-conjugated goat anti-chicken IgY (H+L) cross-adsorbed secondary antibody, Alexa Fluor 488-conjugated donkey anti-rat IgG (H+L) highly cross-adsorbed secondary antibody, and Alexa Fluor 594-conjugated goat anti-rabbit IgG (H+L) highly cross-adsorbed secondary antibody.

A 10 mM oleic acid stock solution was made in 3 mM fatty acid–free BSA (Millipore-Sigma)-PBS. The solution was incubated in 37°C shaking incubator for an hour to completely dissolve oleic acid in the 3 mM BSA-PBS. The stock solution was filtered and stored in -20°C.

### Plasmid construction

The following plasmids were kind gifts: pCMV 3XFLAG-LC3A WT (Addgene plasmid #123089), pCMV 3XFLAG-LC3B WT (Addgene plasmid #123092), pCMV 3XFLAG-LC3C WT (Addgene plasmid #123095), pCMV 3XFLAG-GABARAP WT (Addgene plasmid #123097), pCMV 3XFLAG-GABARAPL1 WT (Addgene plasmid #123100), pCMV 3XFLAG-GABARAPL2 WT (Addgene plasmid #123103) from Robin Ketteler, pSpCas9(BB)-2A-Puro (PX459) V2.0 (Addgene plasmid #62988) from Feng Zhang, and pmScarlet-I_C1 (Addgene plasmids #85044) from Dorus Gadella. pEGFP-N1 and pEGFP-C1 plasmids were purchased from Clontech Laboratories, pSMART-HC-Amp plasmid was purchased from Lucigen. pET28-12XHis-SUMO plasmid was from Xudong Wu.

For AAV constructs, ENTR clones were first generated by inserting PCR fragments encoding either mScarlet-I or mScarlet-I-spartin (380-666 aa) using AgeI and KpnI sites. Both PCR fragments were generated from pCMV-mScarlet-I-spartin with primer sets: 5′-GCACAACCGGTGCCACCATGGTGAG CAAGG-3′ and 5′-CGGGGTACCTCACTTGTACAGCTCGTCCATGCC-3′ (for mScarlet-I), 5′-GCACA ACCGGTGCCACCATGGTGAGCAAGG-3′ and 5′-CGGGGTACCTCATTTATCTTTCTTCTTTGCCT CCTTTACTTCCT-3′ (for mScarlet-i-spartin (380-666 aa)). The AAV constructs were then generated by Gateway LR recombination using a destination vector of the modified pAAV-MCS (*38*) containing a synapsin promoter and a woodchuck hepatitis virus post-transcriptional regulatory element (WPRE).

For other plasmid construction, all PCRs were performed using PfuUltra II Fusion HotStart DNA Polymerase (Agilent Technologies) and restriction enzymes were from New England Biolabs. The synthetic DNAs (gBlock, Integrated DNA Technologies) that were used in this study and cloning strategies of the other plasmids (including primer information) were summarized in **table S1** and **S2**.

### Generation of KI and KO cells with CRISPR/Cas9-mediated genome editing

A spartin-knockout (KO) SUM159 cell line was generated by CRISPR/Cas9 gene editing from a wild-type background (*39*). The sequence 5′-CTCTACAGAATGTACGCACC-3′ was used as a gRNA to direct Cas9 into the exon 2 of the *SPG20/spartin* locus. Cells were selected with 1.5 μg/ml puromycin for 48 h. Genomic DNA of clones showing depletion of spartin protein by immunoblot analysis with the spartin antibody were extracted (DNeasy Blood and Tissue Kit, Qiagen), and the genomic DNA sequence surrounding the target exon of *spartin* was amplified by PCR (sense: 5′-AAATGGAGCAAGAGCCACAAAATGGAG-3′, antisense: 5′-GAGGAGCTTCTGCTGGACAACTTTGTG-3′). PCR products were subcloned into a plasmid (Zero Blunt TOPO PCR Cloning Kit, Thermo Fisher Scientific) to validate the edited region of positive KO clones by sequencing.

To generate N-terminally mScarlet-I-tagged spartin and N-terminally mScarlet-I-tagged PLIN3, SUM159 cells were transfected by FuGENE HD transfection reagent (Promega Corporation) with an individual donor plasmid containing arms with ∼ 800-nucleotide-long homology upstream and downstream of the target site and a gRNA targeting downstream of start codon (for spartin, 5′-CTCTACAGAATGTAC GCACC-3′), and downstream of start codon (for PLIN3, 5′-AGAGACCATGTCTGCCGACG-3′), respectively. The homology arm sequence information of individual donor plasmid (pSMART-mScarlet-I-spartin and pSMART-mScarlet-I-PLIN3) are described in **table S1 and S2**. Cells were selected with 1.5 μg/ml puromycin for 48 h, and single-cell FACS sorting was performed by fluorescent proteins at the flow cytometry core facility (Harvard, Division of Immunology). To validate the insertion of tags, target regions were amplified by PCR and sequenced.

The double KI SUM159 cell line (HaloTag-KI-spartin and mScarlet-I-KI-PLIN3) was generated by sequential clonal generation. The mScarlet-I-KI-PLIN3 cell line was initially generated and used for the generation of double KI cell line. The donor plasmid (pSMART-HaloTag-spartin) is described in **table S1 and S2**.

### Transfection and RNA interference

Transfection of plasmids into SUM159 cells was performed with FuGENE HD transfection reagent (Promega Corporation) ∼24 h prior to imaging.

Specific knockdown of spartin in SUM159 cells was performed by transfection of small interfering RNA (siRNA) duplexes by Lipofectamine RNAiMAX (Life Technologies). The siRNAs for Negative Control (#D-001220-01) and ATGL/PNPLA2 (#L-009003-01-0005) were purchased from Dharmacon.

### Expression and purification of GST-fusion proteins and UBR-HA

GST-fusion proteins and 12xHis-SUMO-UBR-HA were expressed in NiCo21(DE3) competent *E. coli* cells (New England Biolabs). Expression was induced with 0.5 mM IPTG in a 1-liter culture of LB (supplemented with ampicillin) for 20 h at 18°C. Cells were harvested by centrifugation.

GST-fusion protein expressing cells were resuspended in buffer A (50 mM Tris-HCl pH 8.0, 500 mM NaCl, 0.5 mM DTT, and cOmplete Protease Inhibitor Cocktail tablet, EDTA-Free (Millipore Sigma)) and lysed by sonication. The cell debris was removed by centrifugation at 5000 g for 15 min. The supernatant was incubated with 1 ml of Glutathione Sepharose 4B (Cytiva) for 2 h at 4°C. The resins were collected and washed with 30 ml of buffer A and once with 10 ml of buffer (50 mM Tris pH 8.0, 150 mM NaCl). GST-fusion proteins were eluted with 2 ml of elution buffer (50 mM Tris pH 8.0, 150 mM NaCl, 10 mM reduced glutathione).

12xHis-SUMO-UBR-HA expressing cells were resuspended in buffer (50 mM Tris-HCl pH 8.0, 500 mM NaCl, 10 mM imidazole, 0.5 mM DTT, and cOmplete Protease Inhibitor Cocktail tablet, EDTA-Free (Millipore Sigma)). Cells were lysed by sonication. The cell debris was removed by centrifugation at 5000 g for 15 min. The supernatant was incubated with 1 ml of PureCube 100 INDIGO Ni-Agarose (Cube Biotech) for 2 h at 4°C. The resins were then collected and washed with 30 ml of buffer (50 mM Tris pH 8.0, 300 mM NaCl, 25 mM imidazole) and eluted with buffer (50 mM Tris pH 8.0, 300 mM NaCl, 500 mM Imidazole). Once imidazole concentration was reduced to 10 mM by sample concentration and re-dilution in buffer (50 mM Tris pH 8.0, 300 mM NaCl), 12xHis-SUMO tag is removed by SUMO protease (a gift from X. Wu) for overnight at 4°C.

The eluted proteins were concentrated and further purified by size-exclusion chromatography on a Superdex200 column, equilibrated with buffer containing 50 mM Tris pH 8.0, 150 mM NaCl, 1 mM DTT. Peak fractions were pooled and concentrated for *in vitro* assay.

### Immunoprecipitation and immunoblotting

For protein-level analyses (fig. S3, B to D and fig. S5), cells were lysed in 1% SDS lysis buffer (50 mM Tris, pH 8.0, 150 mM NaCl, 1% SDS, and cOmplete Protease Inhibitor Cocktail tablet, EDTA-Free (Millipore Sigma)) with ∼100 units of Benzonase® Nuclease (Millipore-Sigma). After protein concentrations were determined using Pierce™ BCA Protein Assay Kit (Thermo Fisher Scientific), cell lysates were mixed with Laemmli sample buffer and heated for 10 min at 75 °C prior to SDS-PAGE.

For immunoprecipitation in Fig. 2A, cells were lysed in 500 μl of lysis buffer (50 mM Tris-HCl, pH 8, 150 mM NaCl, 1 mM EDTA, 1% Triton X-100, PhosSTOP (Millipore-Sigma), and cOmplete Protease Inhibitor Cocktail tablet, and EDTA-Free (Millipore-Sigma)). After rocking for 20 min at 4°C, the cell lysates were centrifuged at 18,000 *g* for 10 min at 4 °C, and the supernatants were collected. The 400 μl of protein lysates were incubated with 20 μl of anti-FLAG® M2 affinity gel (Millipore-Sigma) for 2 h at 4°C. The bead-bound materials were washed with 500 μl of washing buffer (50 mM Tris-HCl, pH 8, 150 mM NaCl, 1 mM EDTA, and 1% Triton X-100) for three times and were eluted with Laemmli sample buffer for 10 min at 75°C prior to SDS-PAGE.

For immunoblot analyses, all gels were transferred to Immuno-Blot PVDF membranes (Bio-Rad) with 1X Tris/glycine transfer buffer (Bio-Rad) with 20% methanol for 1.5 h at 100 V in a cold room. The membranes were incubated in TBS-T supplemented with 5% non-fat dry milk (Santa Cruz Biotechnology) at room temperature for 20-60 min and subsequently incubated with primary antibodies for overnight in cold room with gentle shaking. Membranes were washed three times in TBS-T for 5 min each and incubated at room temperature for 60 mins with appropriate HRP-conjugated secondary antibodies (Santa Cruz Biotechnology) prior to analysis by chemiluminescence with the SuperSignal West Pico or Dura reagents (Thermo Fisher Scientific).

For Coomassie staining, SDS-PAGE gels were washed with Milli-Q water for 20 mins to get rids of residual SDS, then the gels were incubated with colloidal Coomassie staining buffer (10% ethanol, 0.02% Coomassie brilliant blue G-250, 5% aluminum sulfate-(14–18)-hydrate, 2% ortho-phosphoric acid, 85%) for more than 3 h.

### GST-LC3A pull-down with HEK293T cell extracts

The indicated GST-fusion proteins (3 nM) were equilibrated in 500 μl of assay buffer (50 mM Tris-HCl, pH 8, 150 mM NaCl, 1% Triton X-100, PhosSTOP (Millipore-Sigma), and cOmplete Protease Inhibitor Cocktail tablet, and EDTA-Free (Millipore-Sigma)) and mixed with 10 μl of MagneGST™ Glutathione Particles (Promega) for 1.5 h at 4°C. Meanwhile, the HEK293T cell lysates transiently expressing mScarlet-I-tagged spartin truncation mutants were lysed in 500 μl of lysis buffer (50 mM Tris-HCl, pH 8, 150 mM NaCl, 1 mM EDTA, 1% TritonX-100, PhosSTOP (Millipore-Sigma), and cOmplete Protease Inhibitor Cocktail tablet, and EDTA-Free (Millipore-Sigma)). After rocking for 20 min at 4°C, the cell lysates were centrifuged at 18,000 *g* for 10 min at 4°C, and the supernatants were collected. Subsequently, 500 μg of the cell lysates were mixed with GST-conjugated LC3A for 2 h at 4°C. The resin was washed with wash buffer (50 mM Tris-HCl, pH 8, 150 mM NaCl, and 1% Triton X-100) for three times and were eluted with Laemmli sample buffer for 10 min at 75°C prior to SDS-PAGE.

### In vitro binding assay for GST-LC3A and spartin-UBR

The in vitro binding assay was performed by incubation of the indicated protein combinations (1 μM of each protein) in 300 μl of assay buffer (25 mM Tris-HCl, pH 8, 150 mM NaCl, and 5% glycerol) for 2 h at 4°C. The resin was washed with wash buffer (25 mM Tris-HCl, pH 8, 150 mM NaCl, and 0.1% Triton X-100) for three times and were eluted with Laemmli sample buffer for 10 min at 75°C prior to SDS-PAGE.

### Animals

C57BL/6 wild-type mice were obtained from The Jackson Laboratory (Stock No: 000664). Mice were maintained at the Division of Laboratory Animal Resources (DLAR) facility of Wayne State University. All the procedures were conducted under the guidelines of the respective Institutional Animal Care and Use Committee.

### AAV production

AAVs were generated using AAV-DJ Helper Free system (Cell Biolabs) as reported previously (*38*). An AAV construct was transfected with pAAV-DJ and pHelper (Cell Biolabs) into 293FT cells (ThermoFisher). Transfected cells were lysed, and AAVs were purified with HiTrap Heparin HP columns (Cytiva) as previously described (*40*). The titer of each AAV was evaluated by quantitative PCR.

### Virus injection

Seven- to eight-week-old C57BL/6 mice were anesthetized with 1.5-3.0% isoflurane and placed in a stereotaxic apparatus (Kopf Instruments). Subcutaneous injections of meloxicam (5 mg/kg body weight) were administered for three consecutive days, beginning one day before the operation. The skull was exposed over the M1 motor cortex based on stereotaxic coordinates. Then, 2 μl of AAV (approximately 6 × 10^7^ vg) was injected into each hemisphere of the M1 motor cortex using a glass pipette (tip diameter 5∼8 mm) at a rate of 100 nl/min using a syringe pump (Micro4; World Precision Instruments). The injection site was standardized among animals by using stereotaxic coordinates (ML = ±1.50, AP = +1.00, DV = –1.50 and –1.25) from bregma. At the end of the injections, we waited at least 10 min before retracting the pipette.

### Histology

Nine-to ten-week-old wild-type mice were deeply anesthetized and transcardially perfused with 4% PFA in PBS (pH 7.4). Brains were fixed overnight in 4% PFA, and 50 μm thick sagittal sections were cut on a vibratome (Leica Biosystems) at 4°C. Free-floating sections were washed in PBS and then incubated for 45 min at room temperature with 5% normal goat serum and 0.3% Triton X-100 in PBS. Slices were incubated overnight at room temperature with PBS containing 3% normal goat serum, 0.1% Triton X-100, and primary antibodies: rabbit polyclonal anti-spartin (1:500), chicken polyclonal anti-MAP2 (1:1,000), and rat polyclonal anti-GFAP (1:1,000). Sections were then washed three times in PBS and incubated for 2 h at room temperature with PBS containing 3% normal goat serum and secondary antibodies: Alexa Fluor 594-conjugated goat anti-rabbit IgG, DyLight 488-conjugated goat anti-chicken IgY, and Alexa Fluor 488- conjugated donkey anti-rat IgG. For BODIPY493/503 staining, AAV-injected brains were fixed overnight in 4% PFA and cut into 50 μm thick coronal sections on a vibratome at 4°C. Free-floating sections were washed three times in PBS and then incubated for 10 min at room temperature with 0.3 μM BODIPY493/503 in PBS. After staining, sections were washed three times in PBS and mounted on microscope slides with Vectashield antifade mounting media (Vector Laboratories). Confocal fluorescence images were acquired on a laser scanning confocal microscope (LSM 780; Zeiss) equipped with a 10× (NA 0.3) objective or a 63× (NA 1.4) objective.

### Lipidomic profiling of mouse brain tissues

Twelve days after AAV injection, mScarlet-I-positive regions of the M1 motor cortex were dissected in ice-cold PBS under an epifluorescent microscope (CKX53, Olympus). Dissected tissues were immediately frozen in liquid nitrogen and stored at -80°C until lipid extraction. Excised brain tissues were then homogenized in ice-cold nuclease-free water using Bead Mill homogenizer (VWR). Lipids were extracted from 7.5 mg of tissue homogenate containing 8 μl SPLASH LIPIDOMIX mass spec standard (Avanti Polar Lipids) with the addition of methyl tert-butyl ether (MTBE) (Sigma Aldrich) and methanol (Sigma Aldrich) under 7:2:1.5 (v/v) mixing ratio of MTBE, methanol and tissue homogenate, respectively. Samples were mixed and incubated on a thermo shaker for 60 min at 4°C with 1,000 rpm agitating speed. Following extraction, samples were centrifuged at 10,000 x g for 5 min at 4°C. Organic upper-layer were transferred into fresh microfuge tube and dried using vacuum concentrator (Eppendorf).

Lipids were separated using the Thermo Acclaim C30 reverse-phase column (ID 2.1 x 250 mm, 3 μM pore size) (Thermo Fisher Scientific) connected to a Dionex UltiMate 3000 UHPLC system and a Q-Exactive orbitrap mass spectrometer (Thermo Fisher Scientific). Extracted lipid samples were dissolved in 150 μl solvent containing chloroform/methanol (2:1 v/v). 10 μl of lipids were injected for analysis positive and negative ionization modes, respectively. UHPLC solvents consist of stationary phase 60:40 water/acetonitrile (v/v), 10 mM ammonium formate and 0.1% formic acid, and mobile phase 90:10 isopropanol/acetonitrile (v/v), 10 mM ammonium formate and 0.1% (v/v) formic acid. Lipids were separated over 90 min gradient at 55°C with chromatographic flow-rate of 0.2 mL/min. Mass spectrometer data were converted to mzML format using MSConvert (*41*) prior to analysis using LipidXplorer version 1.2.8.1 (*42*) and duplicate lipids were excluded. Each lipid classes were normalized by total phosphotidylcholine level since total phosphotidylcholine were not significantly changed across the samples. Statistical analyses on lipid abundance were calculated using Prism8.

### Lipid extraction and thin layer chromatography

Cells in 6-well cell-culture plates were pulse-labeled with 500 μM [^14^C]-oleic acid (50 μCi/μmol) for designated time. Cells were washed with ice-cold PBS for two times, then harvested by trypsinization. Cell pellets were collected by centrifugation at 600 g for 5 min and resuspended with 85 μl Milli Q water on ice. For DNA measurement, 5 μl of total resuspension was aliquoted into 95 μl Milli Q water and lysed by freezing-thawing and subsequent 1 h incubation at 37 °C with gentle shaking. DNA samples were prepared with FluoReporter blue fluorometric dsDNA quantification kit (Thermo Fisher Scientific) and measured by TECAN microplate reader (TECAN). For lipid extraction, rest of 80 μl of total lysates were mixed with 300 μl of CHCl3:MeOH (1:2, v/v), vortexed, mixed with 100 μl of CHCl3, vortexed, and mixed with 100 μl of Milli Q water. After centrifugation at 2000 g for 5 min, organic bottom-layers were transferred into fresh microfuge tube and dried under nitrogen stream. Samples were normalized by DNA concentration and separated by TLC with hexane: diethyl ether: acetic acid (80:20:1) solvent system. TLC plate were exposed to phosphor imaging cassette overnight and revealed by Typhoon FLA 7000 phosphor imager. Standard lipids on TLC plate were stained with iodine vapors afterwards.

### Fluorescence microscopy

Cells were plated on 35 mm glass-bottom dishes (MatTek Corp). Imaging was carried out at 37°C approximately 24 h after transfection. For fixed samples, cells were washed with ice-cold PBS twice, followed by incubation with 4% formaldehyde (Polysciences)-PBS for 20 mins at room temperature. After fixation, cells were washed three times with PBS for 5 mins. Where noted, cells were stained with 0.5 μM BODIPY493/503 (Thermo Fisher Scientific) and 1 µg/mL Hoechst 33342 (Thermo Fisher Scientific) approximately 20 mins before imaging.

Spinning-disc confocal microscopy was performed using a Nikon Eclipse Ti inverted microscope equipped with Perfect Focus, a CSU-X1 spinning disk confocal head (Yokogawa), Zyla 4.2 Plus scientific complementary metal-oxide semiconductor (sCMOS) cameras (Andor, Belfast, UK), and controlled by NIS-Elements software (Nikon). To maintain 85% humidity, 37°C and 5% CO2 levels, a stage top chamber was used (Okolab). Images were acquired through a 60× Plan Apo 1.40 NA objective or 100× Plan Apo 1.40 NA objective (Nikon). Image pixel sizes were 0.107 and 0.065 μm, respectively. Blue, green, red, and far-red fluorescence was excited by 405, 488, 560, or 637 nm (solid state; Andor, Andor, Cobolt, Coherent, respectively) lasers. All laser lines shared a quad-pass dichroic beamsplitter (Di01-T405/488/568/647, Semrock). Blue, green, red, and far-red emission was selected with FF01-452/45, FF03-525/50, FF01-607/36, or FF02-685/40 filters (Semrock) respectively, mounted in an external filter wheel. Multicolor images were acquired sequentially.

High-throughput imaging was performed on an IN CELL Analyzer 6000 microscope (GE Healthcare Life Sciences) using a 60× 0.95 NA objective lens. Cells were prepared in 24-well glass-bottom plates and fixed in 4% formaldehyde-PBS at room temperature for 20 mins, washed three times with PBS, and stained with 0.5 μM BODIPY493/503 for LDs and Hoechst 33342 (Thermo Fisher Scientific) for nuclei. 20 images were acquired per well. LD areas and numbers from high-throughput microscopic images were quantified using CellProfiler software (*43*).

### Image Processing and Quantification

All acquired images were processed and prepared for figures using Fiji (*44*). For co-localization analyses of Fig. 3B and 3E, each of the 6-8 cropped images containing cell-peripheral regions (where LDs preferentially co-localize with lysosomes) were manually prepared from 6 different cell images. Then, co-localization between LDs (either marked with LiveDrop or spartin) and lysosomes (marked with LAMP1-mNG) were determined by Pearson’s coefficient analysis using “Coloc 2” tool in Fiji. For LD number quantification shown in fig. S7B, more than 30 cells of each conditions were analyzed from 10 images. Cell outlines were manually drawn, and LDs in BODIPY channel were identified by “Find maxima” tool in Fiji.

### Image analysis was performed in python for the following figures

Figure 1B: For each of the 12 images analyzed (6 images in 2 conditions) a single cell entirely contained within the image was manually outlined with napari (https://github.com/napari/napari). Image intensities were background subtracted and “total spartin” content was calculated as the sum intensity of the spartin channel falling within the manually outlined cell mask. Lipid droplet centroids and radii were detected using the Laplacian of Gaussian (LoG: skimage.feature.blob_log) method from scikit-image(*45*). “LD-localized spartin content” was calculated as the sum intensity of pixels in the spartin channel falling inside of the radii of one of the detected lipid droplets. “Spartin enrichment at LDs” is defined as the ratio of LD-localized spartin to total spartin.

Fig S1D: PLIN3 enrichment at LDs was calculated in the same way as described above for spartin, but for the PLIN3 channel.

Figure 1D: In three-channel images of Spartin, PLIN3, and lipid droplets, lipid droplets were again detected using LoG blob detection, and a binary lipid droplet mask was created using the detected centroids and radii. Pixels from both the spartin and PLIN3 channel were masked using the lipid droplet mask, and the “lipid-droplet-localized” Pearson correlation coefficient between spartin and PLIN3 was calculated using numpy.corrcoef(*46*).

Figure 4I: 3D confocal images were analyzed plane-by-plane (z-step size was significantly larger than the lipid droplets being detected). Cell body boundaries were detected with Cellpose (*47*), using the “ctyo2” pretrained model with a cell diameter of 174 pixels and a cell probability threshold of -1.8. For each image, the “total cell area” was calculated as the total number of pixels falling within a cell body multiplied by the area of a single pixel (0.065 µm pixel size for an area of 0.004225 µm^2^). The coordinates of putative lipid droplets were first detected using LoG as described above. To reduce false positives from non-specific tissue, “true mature” lipid droplets were filtered as follows: In a small image patch around a putative LD coordinate (stained by BODIPY493/503), the correlation between the BODIPY and mScarlet-I channels was measured. Because true mature lipid droplets are generally negatively correlated with mScarlet-I (i.e., the core of matured LD stained with BODIPY cannot be correlated with mScarlet-I- as mScarlet-I-tagged spartin-DN decorates the surface of the LD monolayer), only LDs with a negative correlation coefficient between the LD and mScarlet-I channel were included. Finally, for each image analyzed we report the number of true lipid droplets detected normalized to the total cell body area.

### Statistics

Unless otherwise stated, results are presented as mean ± standard deviation. Statistical analyses of results were performed using GraphPad Prism 8 (for statistical details of each experiment, see figure legends). Statistically significant differences are denoted as follows: *p < 0.05, **p < 0.01, ***p < 0.001, ****p < 0.0001.

---

**Table S1.**
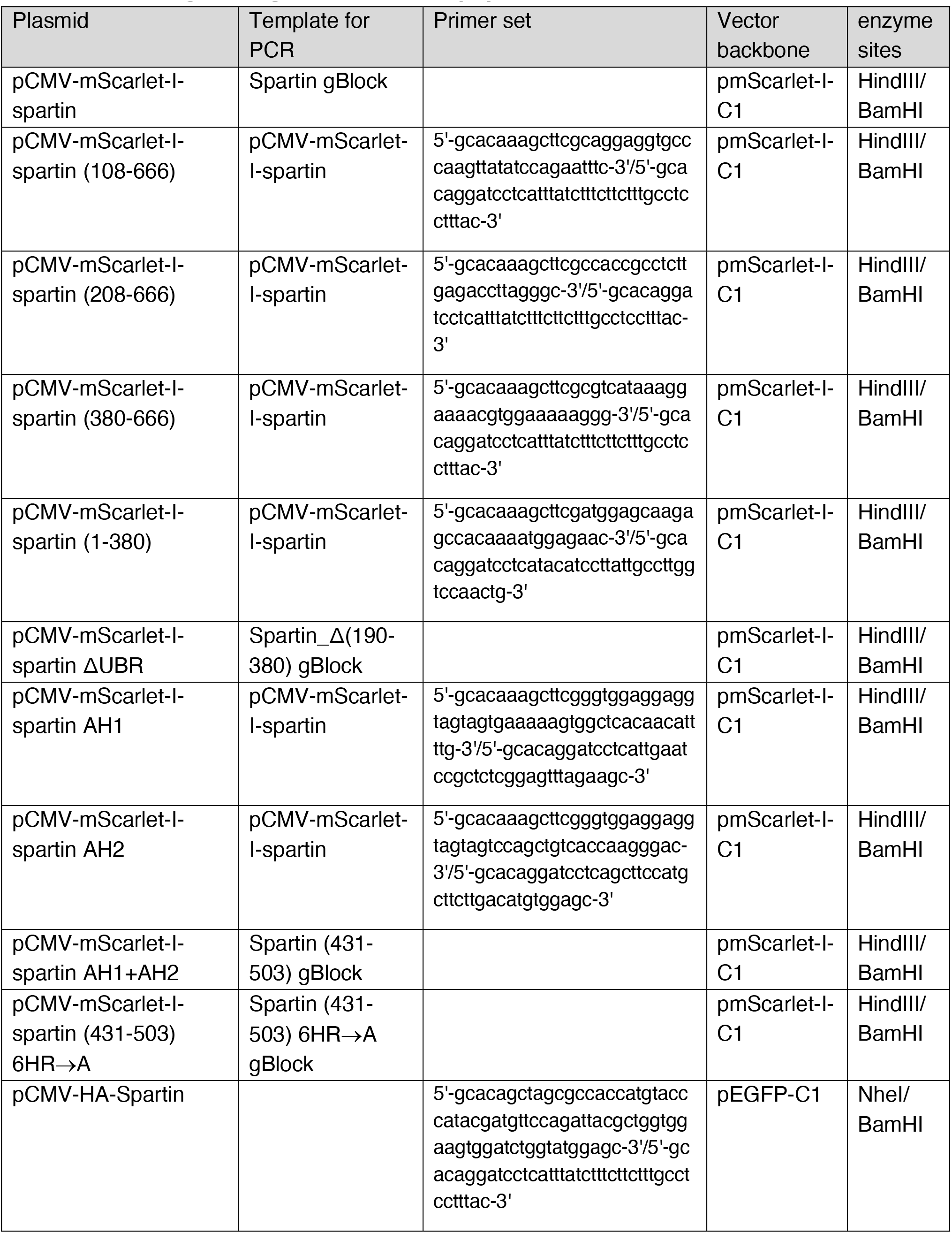
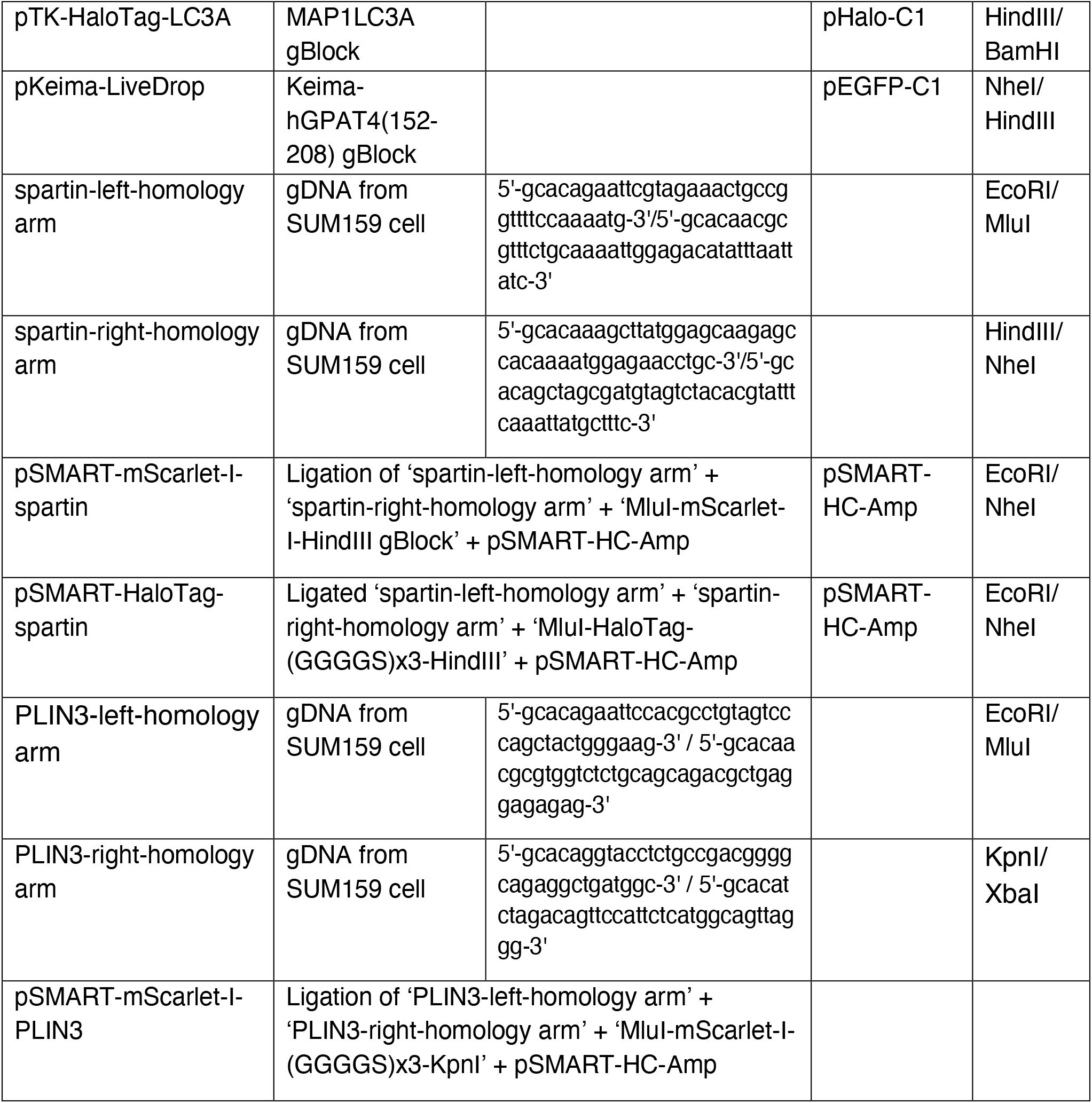
**Cloning strategies used in this paper.**

**Table S2.**
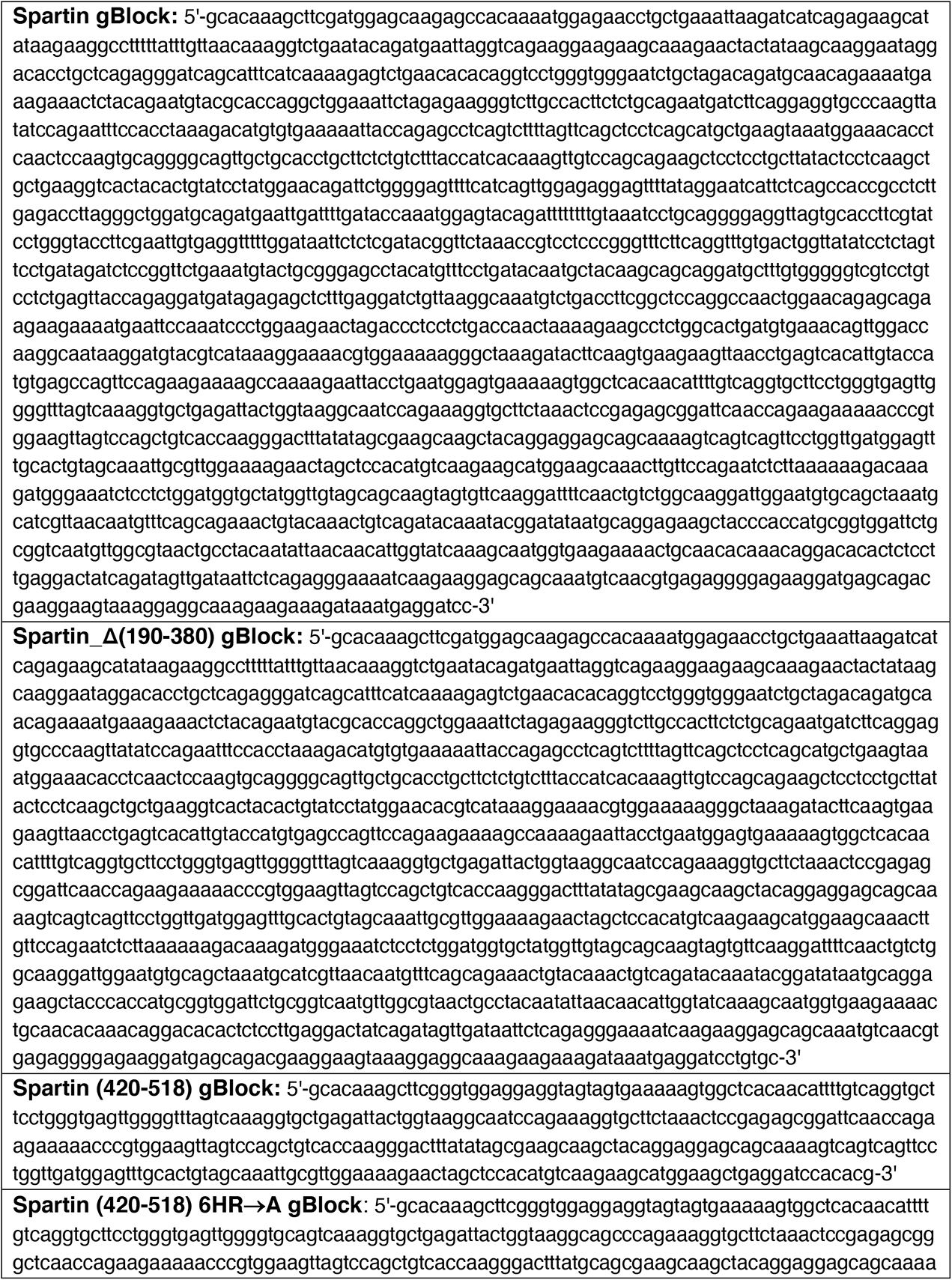
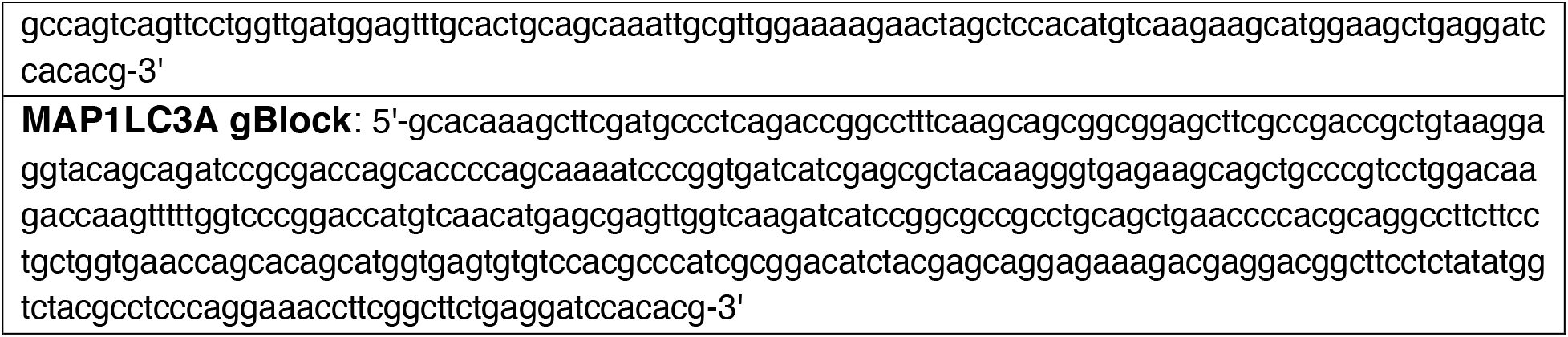
**Synthesis DNAs (gBlock) used in this paper.**

